# Polymer dynamics relates chromosome mixing to temporal changes in biological contact maps

**DOI:** 10.1101/2022.07.20.500905

**Authors:** Gaurav Bajpai, Samuel Safran

## Abstract

Chromosomes are arranged in distinct territories within the nucleus of animal cells. Recent experiments have shown that these territories overlap at their edges, suggesting partial mixing during interphase. Genomewide, biological contact maps in humans and *Drosophila* show only a low degree of contact between different chromosomes; however, the mixing in yeast is considerably higher. Recent theoretical estimates considered topological mixing of chromosomes by polymer reptation, and suggested that the time scale for chromosome mixing is years. This implies that a cell will typically divide before its chromosomes mix by reptation. Here, we use a generic polymer simulation to quantify the dynamics of chromosome mixing over time. We introduce the chromosome mixing index that quantifies the mixing of distinct chromosomes in the nucleus. We find that the chromosome mixing index increases as a power-law with time, and the scaling exponent varies non-monotonically with self-interaction and volume fraction. By comparing the chromosome mixing index with both subdiffusion due to (nontopological) intermingling of chromosomes as well as longer-time reptation, we show that the scaling exponent of the chromosome mixing index is related to intermingling for relatively small chromosome attractions and to reptation for large attractions. The model is extended to realistic biological conditions.

## I. INTRODUCTION

At the nuclear mesoscale, chromosomes can be represented by polymer-like structures in which DNA is tightly packaged along with many histone and non-histone proteins [1] into long chromatin chains. Most eukaryotic cells contain multiple chromosomes within the confines of the cell nucleus. Each individual chromosome condenses when a cell is about to divide (mitosis). In interphase, chromosomes decondense within the nucleus; but instead of mixing, they become organized in distinct regions (territories) of the nucleus [2]. While the chromosomes are organized into distinct territories, overlapping of those territories is also observed in recent studies, suggesting that the different chromosomes are partially mixed at the borders of territories [3–7]. Even if the mixing is far from complete, it can have implications for correlations in gene expression in the regions of the boundaries of the chromosomes that do partially mix within realistic times [8]. The biophysical explanation for why chromosomes do not completely mix during interphase in many organisms is intriguing. Based on reptation theory, that models mixed interphase chromosomes as fully entangled polymers with each chain moving with a snake-like motion (reptation), would predict that the time for mixing scales as the cube of the length of the polymer (*τ* ∝ *N* ^3^ where *τ* is the relaxation/reptation time and *N* is the total length of the polymer) [9–11]. However, mixing can occur without reptation by topological entanglement, just by intermingling of the different polymers. Such motion can be characterized by its mean square displacement (MSD) of a polymer segment which increases with the square root of time as in the Rouse model (MSD ~ *τ* ^1/2^) [9, 12] and with smaller exponents for interacting chains as described below.

Previous studies assumed topological mixing via reptation to estimate the mixing time of chromosomes, and suggested that while the cell cycle time of most animal cells is hours or minutes (for example 24 hours for human cell and 8 minutes for *Drosophila* cell), the mixing time scale for chromosomes is on the order of years (for example 500 years for human cell and 5 years for *Drosophila* cell) [1, 11, 13, 14]. These time scales strongly suggest that the cell will divide long before its chromosomes mix. The results of these studies imply that complete chromosome mixing is not possible during interphase (in real time), but they do not specify the extent of mixing at shorter times. Since even partial mixing implies some correlation in nearby genes (of different chromosomes) it is important to quantify the dynamics of chromosome mixing taking into account the dynamics of chromatin in the nuclear environment that includes the extent of nuclear hydration, protein-induced self-attractions of chromosome segments, and the nuclear lamina. In this paper, we quantify these dynamics using simulations of a generic model of interacting homopolymers and predict the extent and dynamics of mixing as a function of the chromosome segment attraction (*ϵ*) and volume fraction of chain (*ϕ*). In our simulations, the initial condition, motivated by the biological state of separately condensed chromosomes after mitosis is a state where all the chains are separate and confined to the nucleus; as the simulation progresses, the chains begin to mix. This is quantified by our introduction of a single, time-dependent measure of mixing, *α*, easily obtained from the Hi-C (or simulation) chromosomes contact maps. The chromosome mixing index *α* is the ratio of the average number of interchromosomal contacts relative to the average of the intrachromosomal contacts. Our results show that that the polymer chains mix slowly and chain mixing index (*α*) increases with a fractional power of time (*α* ∝ *t*^*β*^ where *β* = 1/4 − 1/8 depends on chain-chain attraction strength and volume fraction of chains). The range of *β* is comparable to the time exponent of the MSD of a chain segment given by Rouse dynamics (when the chain interactions are weak), smaller exponent when chain interactions are relatively stronger, and finally crossing over into the reptation dynamics for strong interactions. Although chromosomes mix slowly, the extent of their mixing is quantifiable and can be accounted for by the mixing index obtains from the contact maps. We estimate the total time for complete chromosome mixing as a function of both *ϕ* and *ϵ*. In addition, we extend our model to include the heterogeneity of the polymers and their lengths to account for eu and heterochromatin and account for the interactions of the chromosomes and the nuclear lamina, to better represent the situation of the *Drosophila* chromosome. We find that the interaction with the lamina further impedes chromosome mixing.

Chromosome conformation capture (3C) based Hi-C experiments can be used to quantify chromosome mixing in different cells. Hi-C experiments measure genome-wide contacts over a population of cells [15]. Human and *Drosophila* Hi-C maps show relatively little interchromosomal (non-diagonal) contacts relative to the number intra-chromosomal (diagonal) contacts, but inter-chromosomal contacts obtained from yeast Hi-C maps are significantly higher [15–17]. A recent single cell, Hi-C study in mammalian cells estimates an approximately 5-10% mixing frequency between chromosomes [18]. Our research has led us to compare the contact maps and defines a single, averaged parameter (the chromosome mixing index *α*) that quantifies the extent of chromosome mixing from both from Hi-C data and simulations. We begin our study with a generic polymer model to understand the physics of chromosome mixing, and then we add other aspects of the nuclear biology one by one, to ascertain what is generic and what is more particular in making the model more specific to *Drosophila* genome. The chromosome mixing index is equal to ratio of the averaged (over many time steps of a simulation, corresponding to short-time or ensemble averages of cells) sum of interchromosomal contacts and and the averaged sum of intrachromosomal contacts. The different chromosomes mix as interphase proceeds in time, and we find that the chromosome mixing index scales with time (*α* ~ *t*^*β*^), increasing in a power-law fashion; the scaling exponent exponent *β* varies in a non-monotonic manner as a function of the chromosome volume fraction and interactions. The main message of our paper is that one can account in a relatively simple manner, and encapsulate in one time-dependent parameter, the extent chromosome mixing from the complex Hi-C data.

In addition to the mixing of different chains (chromosomes), we also the mixing within a single chain by calculating from our simulations, the contact probability, *P* (*s*) as a function of the bead separation along the chain contour, *s*. We find that *P* (*s*) ∝ *s*^*−γ*^, where *γ* is the exponent whose value indicates (when compared with the mixing analysis of the simulations) whether subdomains within a single chain mix or not. It has been reported that contact probability scaling exponent (*γ*) for an unconfined, random-walk, phantom chain is *γ* = 1.5 and for a self-avoiding chain is *γ* = 2.1 [19, 20]. For a collapsed chain, scaling exponent *γ* ≈ 0 which indicates an equilibrium globule, with a high degree of mixing of distant segments along the chain, while *γ* = 1 indicates a fractal globule, is characterized by very little mixing [15, 21, 22] of distant segments. An exponent of *γ* = 0.5 was deduced from the Hi-C experiments of a mitotic chromosome, with a computational polymer dynamics model suggesting intermediate mixing [22, 23]. Furthermore, *γ* = 0.75 observed from Hi-C contact maps of interphase chromosomes and compared with a computational polymer dynamics model suggests very slow mixing (glassy dynamics) [19, 24]. Also, recent studies suggest that chromosome X in mammalian cells forms a glassy globule with a scaling exponent of *γ* = 0.72 ± 0.2 [25]. We note that all these are structural and not dynamical characterizations. In our simulations, in addition to the dynamics of mixing of different chains, we relate to these structural measures of mixing within a single chain by calculating the scaling exponent *γ*. The contact probability provides information about the mixing of different sub-domains within each chain. Since the attraction within and among chains are the same, similar trends are also seen for the for mixing of different chains (If different regions of one chain are mixed due to the attraction, then different chains will also mix in a similar manner). We do this in the context of averaging the contact probabilities of four chains and shown that the long-time steady-state value of *γ* ranges between 0 (equilibrium mixing) and 1 (almost no mixing).

### A. Biophysical properties of chromatin at the mesoscale

Chromatin (interphase chromosome) has been studied theoretically using models of polymers in both good and poor solvents [22, 26–28]. DNA is negatively charged and wraps around the oppositely charged histone octamer proteins; however, there also regions of non-histone-associated, linker DNA and chromatin still has a net negative charge [29]. Due to electrostatic repulsion of DNA and excluded volume, chromatin had often been considered as a self-avoiding polymer in a good solvent. However, there are also many compelling reasons for treating chromatin (as opposed to DNA) as a self-attractive polymer in poor solvents. The chromatin histone tails that are positively charged attract the negatively charged DNA linkers (in regions that can be far along the contour length of the chain, but close in three-dimensional space), leading to chromatin self-attraction and water acting as a poor solvent [29–34]. In addition, the HP1 chromatin-binding proteins are phase separated within the nucleus and may contribute to (hetero)chromatin condensation [35, 36]. Other studies have demonstrated the presence of a gel-like organization of chromatin that is conducive to self attraction [37–39].

In this paper, we analyze the mixing dynamics of polymer chains (as models of chromosomes) by considering the polymer in both good and poor solvents by varying the strength (*ϵ*) of a standard Lennard-Jones potential (see the methods section IV A) that acts between the beads of the chain. If chromatin would behave a polymer in a good solvent (random walk polymer, *R*_*g*_ ~ *N* ^1/2^ and self-avoiding polymer, *R*_*g*_ ~ *N* ^3/5^), its radius of gyration would typically exceed the diameter of the nucleus [40]; the nuclear envelope would then confine the polymer and prevent it from swelling. It is estimated that chromosomes take up 15 to 60% of the nucleus volume in *Drosophila* and 0.4 to 25% in humans [27, 41–43]. The different hydration of nuclei in different circumstances and organisms means that one must consider how chromosome volume fraction influences the dynamics of chromosome mixing. The polymers in our simulations (whether or in good or poor solvent) are confined and we study this effect by varying the volume fraction of chromosomes from *ϕ* = 0.001 to 0.6. Within this range, the confinement diameter can be either smaller or larger than the radius of gyration of an equivalent random walk, allowing polymers mix with or without constraint.

In addition to the effects of hydration (chromatin volume fraction) and interactions, an individual chromosome is not a homogeneous chain (homopolymer). The chromosome is inhomogeneous even at the mesoscale since it contains both open (euchromatin) and compact (heterochromatin) domains [44]. We have therefore expanded the simulations of generic homopolymers to quantify how the heterogeneous nature of chromatin can prevent chromosomes from mixing. Our simulations of each chromosome as a (non-periodic) block copolymer containing both open and compact regions show that chromosome mixing dynamics is faster in the more open euchromatin regions and slower in the more dense heterochromatin regions.

Along with chromosomes, the nucleus also contains the (mostly aqueous) nucleoplasm, nuclear lamina, and chromosome-binding proteins. The nuclear lamina, comprises lamin proteins and lies on the inner surface of the nucleus (nuclear envelope). Its binding to the lamin-associated domains (LAD) of chromatin has been shown to slow down chromosome dynamics [45, 46]. In Section II E, we show from our simulations that that binding of LAD to the nuclear lamina leads to increased chromosome lamina contact that then reduces inter-chromosome contact. Recent experimental study suggests that unlike fixed cells in which the chromosomes fill the nucleus fairly uniformly, in live *Drosophila*, the chromosomes are organized near the lamina layer of the nuclear envelope [41]. Non-uniform, mesoscale distribution of chromatin had been previously discussed in the single-cell context in [47, 48]. In the live fly experiments [41], overexpression of lamin C results in a shift from peripheral to central organization of chromosomes. These observations suggest that peripheral and central organizations are related to the hydration of the live organism nuclei and that dehydrated nuclei (included by cell fixation and/or spreading on slides) can give only conventional organization in which chromosomes fill the nucleus. Theoretical analyses of the experimental work suggest that chromatin acts as a polymer in poor solvent (due to chromatin self-attraction) and that the various types of mesoscale organization are functions of hydration, chromosome-chromosome interactions, and chromosome-lamina interaction [26, 49].

## II. RESULTS

### A. Definition of the chromosome mixing index and its limiting values

The typical contact map of a genome shows intrachromosomal contacts along the diagonal and interchromosomal contacts along the non-diagonal (see Fig. 1 (a)). The Hi-C contact maps of the human and fly genomes shows that non-diagonal contacts are relatively sparse while in the yeast genome, there are relatively more off-diagonal contacts [15, 50]. We analyze these genome contact maps to study how chromosomes mix, and introduce a single, physical measure that we term the chromosome mixing index (*α*), which quantifies in a coarse-grained manner, the extent of chromosome mixing. The chromosome mixing index defined as the ratio of the total number of interchromosomal contacts and the total intrachromosomal contacts. In the methods section IV E, we theoretically show that the ideal limit of the chromosome mixing index is equal to *n* − 1, where *n* is the number of chromosomes within the nucleus, e.g., the maximum value of chromosome mixing index is 3 for a genome with four chromosomes of the same size. In general *α < n*; the value of chromosome mixing indicates the actual mixing of the chromosomes. In order to allow for the maximal amount of chain (chromosome) mixing from simulation, we simulated phantom chains (non-interacting chains, *ϵ* = 0), which mix quickly as there is no crowding effect due to the lack of volume exclusion (see Fig. 1 (b)).

**FIG. 1:**
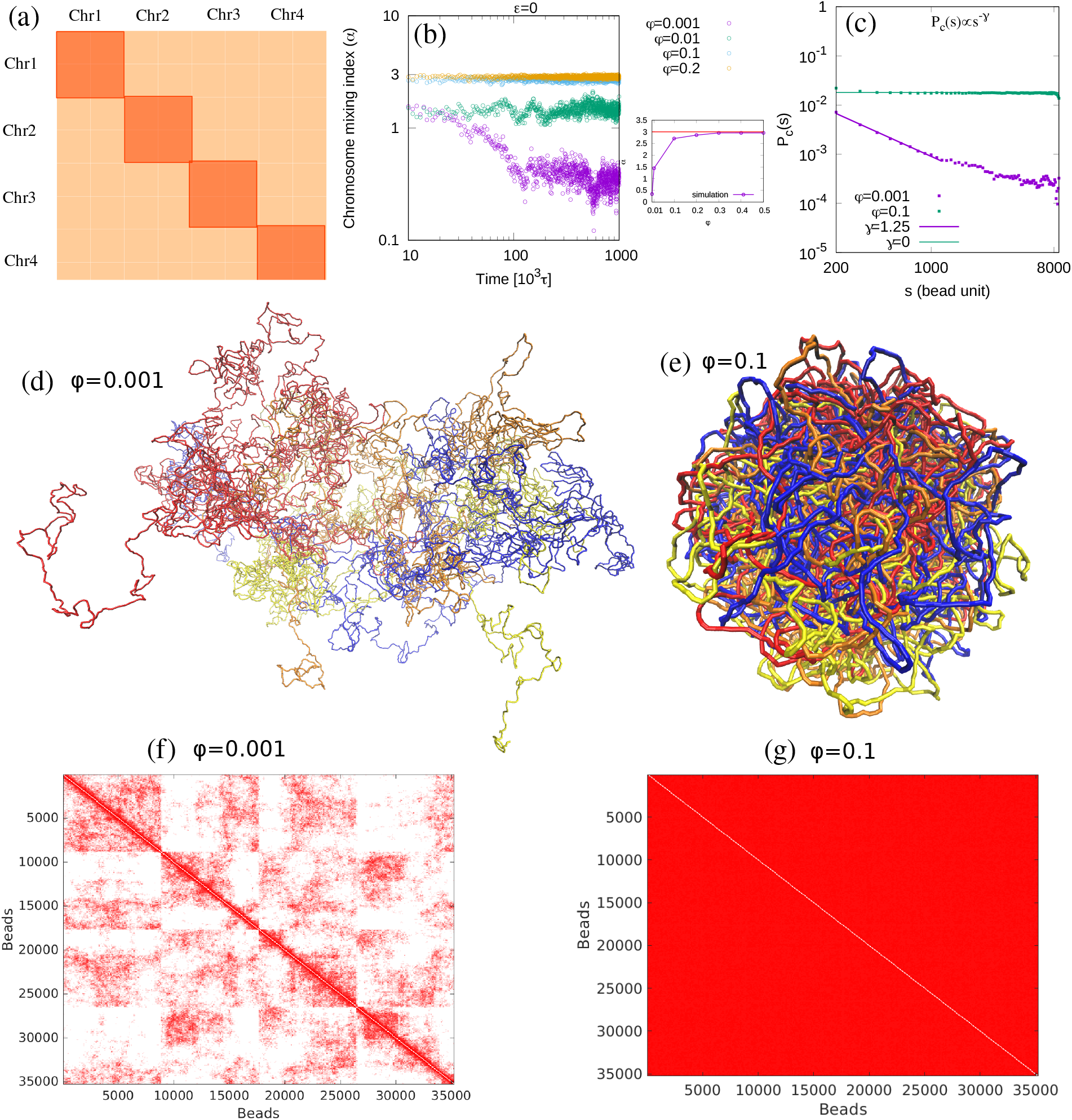
(a) A schematic diagram representing the contact map of a genome consisting of 4 chromosomes. Intra- and interchromosomal contacts are shown by dark and light orange colors, respectively. (b) The evolution of the chromosome mixing index, *α*, over time indicates that phantom chains quickly achieve equilibrium. Right panel of (b) is the time-average of the chromosome mixing index (*α*) as a function of the chain volume fraction (*ϕ*) in the nucleus. For relatively large volume fractions *α* is approximately equal to three (the maximal value of *α* is 3 for this case of 4 chains) which indicates that phantom chains achieve maximal mixing in small confinement volumes. The chromosome mixing index is smaller than 3 for smaller volume fractions because the larger confinement volume (smaller volume fraction of chains), means that any two beads (even within the same chain) are less likely to be “in contact”. (c) The average contact probability *P*_*c*_(*s*) within a single chain, averaged 4 phantom chains as a function of the bead separation distance *s* with the scaling exponent *γ* defined by *P*_*c*_(*s*) ∝ *s*^*−γ*^. (d) Long time snapshot (*t* = 10^6^*τ* time steps) of simulations of 4 phantom chains for a low volume fraction of chains *ϕ* = 0.001, shows that the different chains are not mixed, since they can diffuse away from each other. (e) Late time snapshot (*t* = 10^6^*τ* time steps) of simulations of 4 phantom chains for a relatively large volume fraction *ϕ* = 0.1, shows that the chains are mixed. Note that in Figs. (d) and (e), each color represents a different chain corresponding to a different chromosome. (f) and (g) Contact maps calculated from phantom chain simulations for volume fraction *ϕ* = 0.001 and *ϕ* = 0.1 respectively.

We varied the volume fraction *ϕ* of chains (representing the chromosomes) within the nucleus, and calculated the time-averaged chain mixing index (*α*) as shown in right panel of Fig. 1 (b). The figure shows that phantom chains mix rapidly and attain the maximal value of chromosome mixing index (*α* ≤ 3) for a system with 4 chains, confined to a relatively small volume (a large volume fraction). For larger nuclear volumes (smaller volume fractions of chains), the chromosome mixing index is less than 3 because each chromosome freely diffuses across a large portion of the nucleus and is thereby unlikely to be mixed. We can easily understand this by comparing the radius of gyration and the diameter of confinement. The chromosome mixing index of phantom chains (with radius of gyration *R*_*g*_) reaches its maximal value for relatively small confinement volumes *R*_*c*_, *R*_*g*_ ≫ *R*_*c*_ but has a small value when *R*_*g*_ ≪ *R*_*c*_. In the simulations shown in Fig. A.1(c) (also see the methods section IV A), *R*_*g*_ = 90 *σ* where *σ* is the bead diameter for *ϵ* = 0; different values of *R*_*c*_ correspond to different volume fractions, *ϕ*. Note that for phantom chains, we compare confinement with the radius of gyration corresponding to the length of a single phantom chain rather than the total length of all the chains since the various phantom chains do not interact.

We have also calculated the contact probability from our phantom chain simulations in order to understand how the subdomains of each chain interact (see Fig. 1 (c)). Since we have 4 chains in our simulation, we calculated the contact probabilities of each chain and then took the average, so the final plot shows the average contact probability of 4 chains (see details in methods section IV F). We fit the contact probabilities, *P*_*c*_ as a function of the distance along the contour length of two beads (<u>that belong to the same chain</u>), *s*, with a power law: *P*_*c*_(*s*) ~ *s*^*−γ*^. We then found the scaling exponent *γ* for the range 200 ≤ *s* ≤ 1000 in bead units which is equivalent to the range of 1 ≤ *s* ≤ 5 in mega basepairs. For *ϕ* = 0.1, the scaling exponent (for the large values of *s* of beads near the confinement surface) is *γ* = 0, which indicates that chromosomes form an equilibrium globule [22, 23]. However, for *ϕ* = 0.001, the contact probability of two beads separated by a contour length *s* is characterized by an exponent *γ* = 1.25, which is roughly equivalent to that of an unconfined phantom chain where the Gaussian statistics (and our simulations) result in a scaling exponent *γ* = 3/2 (see Fig. A.1(d)). This result also shows that the scaling exponent, *γ*, (for monomers far apart on the contour of the same chain) of phantom chains decreases with increasing chromosome volume fraction, *ϕ*. The snapshots of the simulation in Figs. 1 (d) and (e) show that the phantom chains do not mix for larger confinement radii (smaller volume fraction *ϕ* = 0.001) but are completely mixed for confinement radii (larger volume fraction *ϕ* = 0.1). Figs. 1 (f) and (g) are contact maps calculated for cases (d) and (e) respectively, obtained by counting beads whose centers are closer than 1.5*σ*, where *σ* is the bead diameter; for typical chromosomal bead, this corresponds to chromosomal beads closer than 45 nm being defined as “in contact”. The contact maps are obtained by averaging over the last 500 snapshots (see details in methods section IV F). The contact maps calculated from the simulation of the phantom chains in Figs. 1 (f) and (g) show a heterogeneous probability distribution for larger confinement and a more homogeneous probability distribution for smaller confinement. From the results, it is clear that the phantom chains mix rapidly in a smaller confinement (when *R*_*g*_ *>* 2*R*_*c*_) and reach the value of chromosome mixing index (*α* ≈ 3 for 4 chains). Additionally, we show in Fig. A.2 that disconnected beads in small confinements (large volume fraction of beads) achieve maximal mixing index for non-zero attraction strengths (*ϵ* = 0.25). This means our analytical argument for the maximal mixing index *β*, are independent of attraction strengths at equilibrium. In the following results, we consider interactions (both excluded volume and attractions) among the beads and calculate the chromosome mixing index by changing the chromosome-chromosome attraction strength *ϵ* and the chromosome volume fraction *ϕ* in our simulations (see the model for the definition of *ϕ* and *ϵ* in methods section IV A).

### B. Effect of chromosome-chromosome attraction on chromosome mixing dynamics

After understanding the results of our simulations of mixing of phantom chains, we next investigated the (more realistic) effects of the interactions of the beads and its effect on the mixing of different chain, minimizing the effect of confinement. We thus performed the simulations in a relatively large confinement volume – larger than the radius of gyration of an individual chain. In particular, we took *R*_*g*_ *<* 2*R*_*c*_, where *R*_*c*_ is the radius of confinement and *R*_*g*_ is the radius of gyration of an unconfined random-walk chain of 4*M* beads (*M* is the total number of beads in one chain and we took *M* = 8810). We simulated chains whose volume fraction in the confinement sphere was *ϕ* = 0.001 and varied the strength of chromosome-chromosome interaction, *ϵ* (see the model in Methods section IV A and simulation parameters in Table 1 in section IV D). To better understand how the attraction strength influences the condensation of a single-chain, we first simulated a single chain without confinement and calculated the radius of gyration as a function of the attraction of the beads (see Appendix A and Fig. A.1). In the Fig. A.3 we also discuss the analytical virial coefficient [51] for the LJ potential which we compare with the value obtained from the simulations at which the chains collapse due to the attractive interactions between the beads.

**TABLE I:** We used reduced unit during the simulation.

| Parameter | Description | reduced unit | SI unit |
| --- | --- | --- | --- |
| T | Temperature | 1.0 | 310 K |
| $k_B T$ | Thermal energy | 1.0 | $4.28 \times 10^{-21}$ J |
| $\epsilon$ | LJ parameter | 0 – 1.0 $k_B T$ | |
| $\sigma$ | Bead diameter | 1.0 | 30 nm |
| $R_c$ | Confinement radius | 20 – 164 $\sigma$ | |
| $k_{\text{spring}}$ | Spring constant | 100 $k_B T / \sigma^2$ | 0.0005 Jm<sup>-2</sup> |
| $l_p$ | Persistence length | 5.0 $\sigma$ | |
| $k_{\text{bend}}$ | Bending stiffness | 5.0 $k_B T$ | $21.4 \times 10^{-21}$ J |
| $\tau$ | Brownian time | $3\pi\eta\sigma^3 / k_B T$ | 60 μs |
| $\Gamma$ | Damping coefficient | 10.0 $\tau$ | |
| $\Delta t$ | Time step | 0.01 $\tau$ | |

For a given value of *ϵ*, computation from the simulations of the radius of gyration of the chain indicates the range of attraction strengths for which the chain is open and those where the chain is collapsed. For *ϵ* < 0.4 k_B_T, an isolated chain is open while for *ϵ* ≥ 0.4 k_B_T, an isolated chain is collapsed. *ϵ*_*c*_ = 0.4 k_B_T is the critical attraction strength at which the chain collapses. Note that for chains with a persistence length of one bead, the critical attraction strength was *ϵ*_*c*_ = 0.3 k_B_T (see Fig. A.1 (b),(c)) which is very close to the value analytically predicted by the vanishing of the second virial coefficient for the LJ potential (see Fig. A.3). We also calculated the mean square displacement (MSD) as a function of time for different *ϵ* values to understand the dynamics of attractively interacting beads (see Appendix A and Figs. A.1 (e),(f)). For weak attraction strengths (still an open and not collapsed chain), *ϵ* = 0 and 0.25, fits of the simulations are close to scaling of MSD ~ *τ* ^1/2^. A scaling exponent of 1/2 is MSD result for a Rouse chain [12] in which interactions are neglected so that the simulations values smaller than ^1^ for these values of *ϵ* indicate the role of attractions that “slow down” (in a scaling sense) the “random walk” of the chain [52]. However, for large attraction strengths, *ϵ* = 0.5, 0.75 and 1, the simulations can be fit with values close to the scaling law MSD ~ *τ* ^1/4^; the value of ^1^ is indeed the MSD scaling exponent appropriate to polymer melts (reptation [9]). This is reasonable, since for large attraction strengths, the chains are very condensed and close to a melt state where the dynamics are slow since they involve reptation of the chain end and not diffusion of the chain in directions perpendicular to its local tangent (see Table 2 in section IV G for variation in value of scaling exponent of MSD as a function of volume fraction and interaction).

To analyze the role of interactions in mixing, we simulated four chains as functions of *ϕ*, starting with the initial condition for which all the chains are separated (see the methods section IV C for the biological significance of this condition). As the simulation proceeds, the chains begin to mix so that intra-chain contacts decrease and inter-chain contacts increase with time. This reflects the non-equilibrium nature of chromosome mixing dynamics. In Fig. 2 (a), we show snapshots of the simulations for different values of the attraction strength *ϵ* (see Fig. A.4 for snapshots of the case where the bead persistence length is unity, *l*_*p*_ = 1 bead). When the attraction strength of the chains is less than the critical strength (*ϵ < ϵ*_*c*_) for collapse, the chains are open and do not mix in a large confinement volume, because two beads (even in the same chain) are unlikely to be “in contact”. At larger attraction strengths (*ϵ* = 1 k_B_T), each chain is collapsed but rarely mixes with the other because the kinetics slow down significantly as the time required to break the bond is *e*^(*zϵ /k*^*B*^*T*)^, where *z* is the number of nearest neighbors (not counting the two nearest neighbors to which each beads is bonded). In addition, the concentration of the chain is relatively larger in the collapsed state, which may make the chain kinetics more “jammed” for the condensed states at large values of the attraction. We find that the chromosome mixing index as a function of time increases with a power law: *α* ∝ *t*^*β*^, where *β* is an exponent whose value characterizes the mixing rate (see Fig. 2 (b)). In the right panel of Fig. 2 (b), we see that the value of the exponent first increases and then decreases as the attraction is increased.

**FIG. 2:**
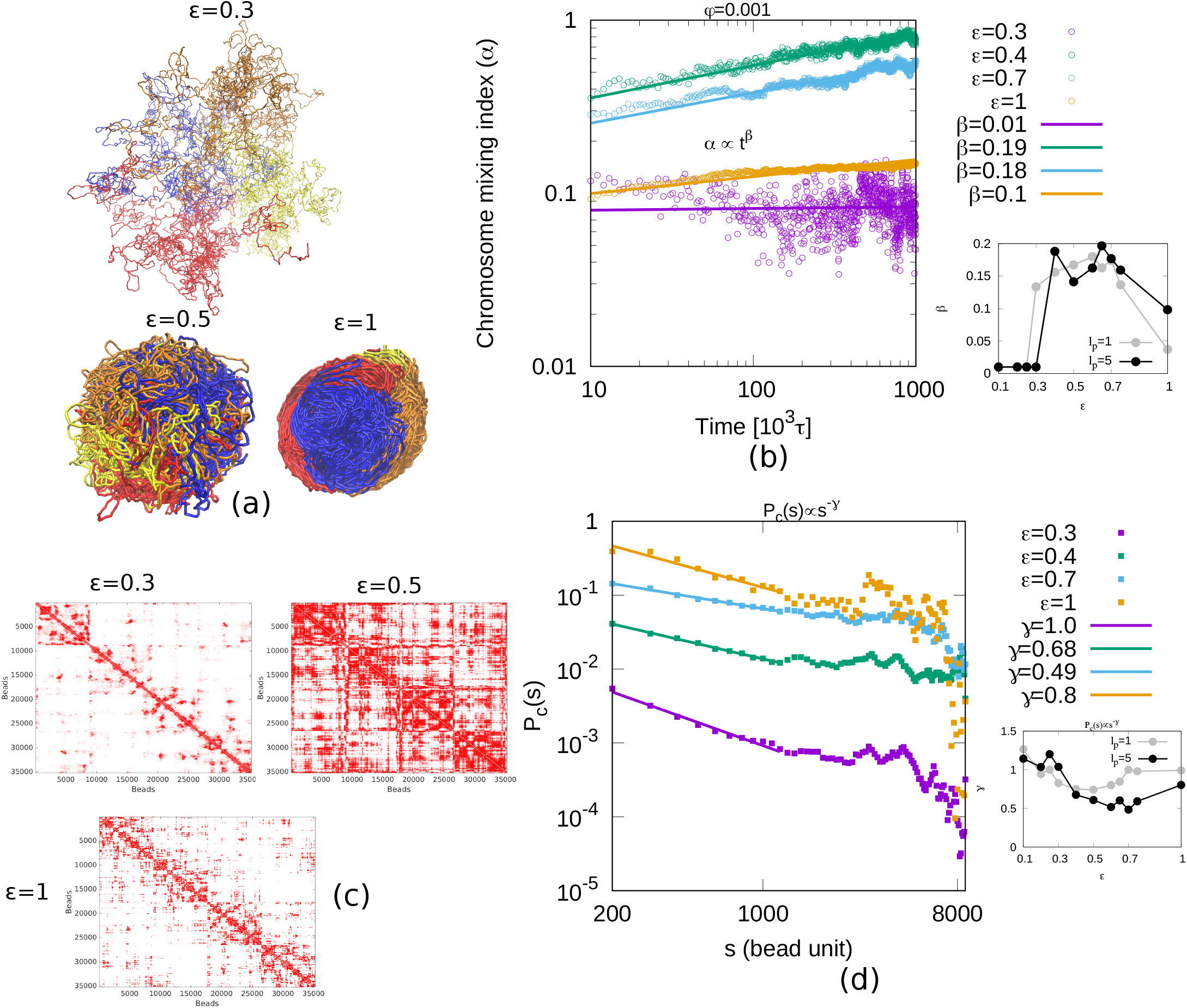
(a) Simulation snapshots at long times (*t* = 10^6^*τ*) shown for different attraction strengths, *ϵ*. Each color represents a different chain corresponding to a different chromosome. For *ϵ* = 0.3, the chains are open and not mixed. For *ϵ* = 0.5, the chains are collapsed and mixed slowly. For *ϵ* = 1, each chain is collapsed but rarely mix with the others. (b) The chromosome mixing index as a function of time increases as a power-law: *α* ∝ *t*^*β*^ where *β* is an exponent, whose value characterizes the rate of mixing. From right panel of (b), we find that the value of the exponent first increases and then decreases as the attraction increases. (c) Contact maps from the simulations calculated by averaging the last 500 frames are shown for different attraction strengths (*ϵ*); in all these simulations, the volume fraction of chains is *ϕ* = 0.001. (d) Average contact probability (*P*_*c*_(*s*)) calculated within each chain is normalized frequency of contact between beads separation distance within a single chain(*s*). Contact probability shows power law relation *P*_*c*_(*s*) ∝ *s*^*−γ*^ where *γ* is exponent. Right panel of (d) is scaling exponent (*γ*) for the contact probability within a single chain as a function of self-attraction strength, *ϵ* for persistence lengths *l*_*p*_ = 1 (gray color) and *l*_*p*_ = 5 (black color).

We calculated the contact maps by averaging the last 500 frames for different attraction strengths (*ϵ*). This allows us to present a time-averaged view of the regions of high or low chromosome mixing (see Fig. 2 (c)). The contact probability (*P*_*c*_(*s*)) calculated from the contact map is the normalized frequency of contact between beads spaced within the same chain, where *s* is their separation along the contour length. We see that the probability obeys power-law scaling with *P*_*c*_(*s*) ∝ *s*^*−γ*^, where *γ* is the exponent (see Fig. 2 (d)). By calculating the contact probability from simulation results, we can find the contact probability scaling exponent for both an open and a collapsed chain. We first calculated the contact probability scaling exponent of a single, unconfined chain (for persistence lengths of both 1 and 5 beads) for various values of the LJ attraction strength, *ϵ* (see Fig. A.1 (d)). The results show that for open chains (*ϵ < ϵ*_*c*_) the contact probability scaling exponent is *γ >* 1 while for collapsed chains the contact probability scaling exponent is *γ* ≤ 1. Furthermore, for the collapsed chains, the scaling exponent 0 *< γ <* 1 shows an intermediate level degree mixing that is a non-equilibrium state between mixing and no mixing. In order to see contact probability behavior across several (four) chromosomes, we calculated contact probability from each chromosome and plotted the average in Fig. 2 (d). In right panel of Fig. 2 (d), we calculated the scaling exponent (*γ*) as a function of attraction strength (*ϵ*) which shows similar trends like single chromosome in which *γ >* 1 for open chains and *γ* ≤ 1 for collapse chain. Collapsed chain shows an intermediate level mixing for 0 *< γ <* 1. We have also shown simulation results of mixing of four chains in relatively small confinement (see Fig. A.5 for simulation snapshots, mixing index, and contact probability. Also see Fig. A.6 and Table IV G for MSD).

### C. Effect of chain volume fraction on mixing dynamics

To examine the effect of spherical confinement (chain volume fraction), we considered open chains (where the attraction is too small to cause collapse) and simulated weakly attractive chains (*ϵ* = 0.25 k_B_T) as a function of *ϕ*. In Fig. 3 (a), we present snapshots of simulations for different volume fractions of chains. For *ϕ* = 0.001, corresponding to large confinement with a relatively small attraction strength (*ϵ* = 0.25), the chains are separated. For *ϕ* = 0.2, the chromosomes begin to mix until each chromosome approximately occupies the entire confinement volume. For *ϕ* = 0.6, the smaller volume restricts the motions of the highly confined (and thus condensed) chains. The nematic ordering observed at this high volume fraction is related to the comparable size of the confinement radius and the persistence length of 5 beads for this simulations, which gives a structure similar to mitotic chromosomes. In the Fig. A.7 we discuss this further and show the structure for a persistence length of one bead where separation, but almost no nematic ordering, is observed. In Fig. 3 (b), the chromosome mixing index as a function of time shows a power-law increase with the exponent *β* (*α* ∝ *t*^*β*^) for various volume fractions of chains. The simulation results give values of *β* that range from 0.001 to 0.25, depending on the volume fraction (confinement). In right panel of Fig. 3 (b), the scaling exponent of the mixing index is plotted as a function of chain volume fraction, which shows that the exponent initially increases and then decreases with chain volume fraction. These power laws suggests that the mixing dynamics is very slow for very large and very small volume fractions (small and large confinement volumes, respectively). In Fig. 3 (c), contact maps obtained from the simulations are calculated by averaging the last 500 frames, and are shown for the different volume fractions, *ϕ*. For *ϕ* = 0.001, the contact map shows fewer intra- and inter-chain contacts which implies that the chromosome chains are relatively open and are hardly mixed. For *ϕ* = 0.2, the contact map shows relatively more intra- and inter-chain contacts which implies that the chromosome chains are both relatively collapsed and mixed. For *ϕ* = 0.6, the contact map shows relatively fewer intra-chain contacts which implies that the dynamics required to cause mixing is impeded; there are also fewer inter-chain contacts which implies relatively little mixing. In Fig. 3 (d), we have plotted the mean contact probability as a function of the bead separation distance, *s*, within a single chain, calculated for the situation of 4 confined chains as the chain volume fraction (*ϕ*) is varied. We find a power law behavior, with *P*_*c*_(*s*) ∝ *s*^*−γ*^ with *γ* exponent. In a large confinement volume (or small chain volume fraction *ϕ* = 0.001), all the chains are open and the contact probability scaling exponent is *γ >* 1. Chains collapse for relatively small confinement (or high chain volume fraction *ϕ* ≥ 0.01), and the scaling exponent is *γ <* 1.

**FIG. 3:**
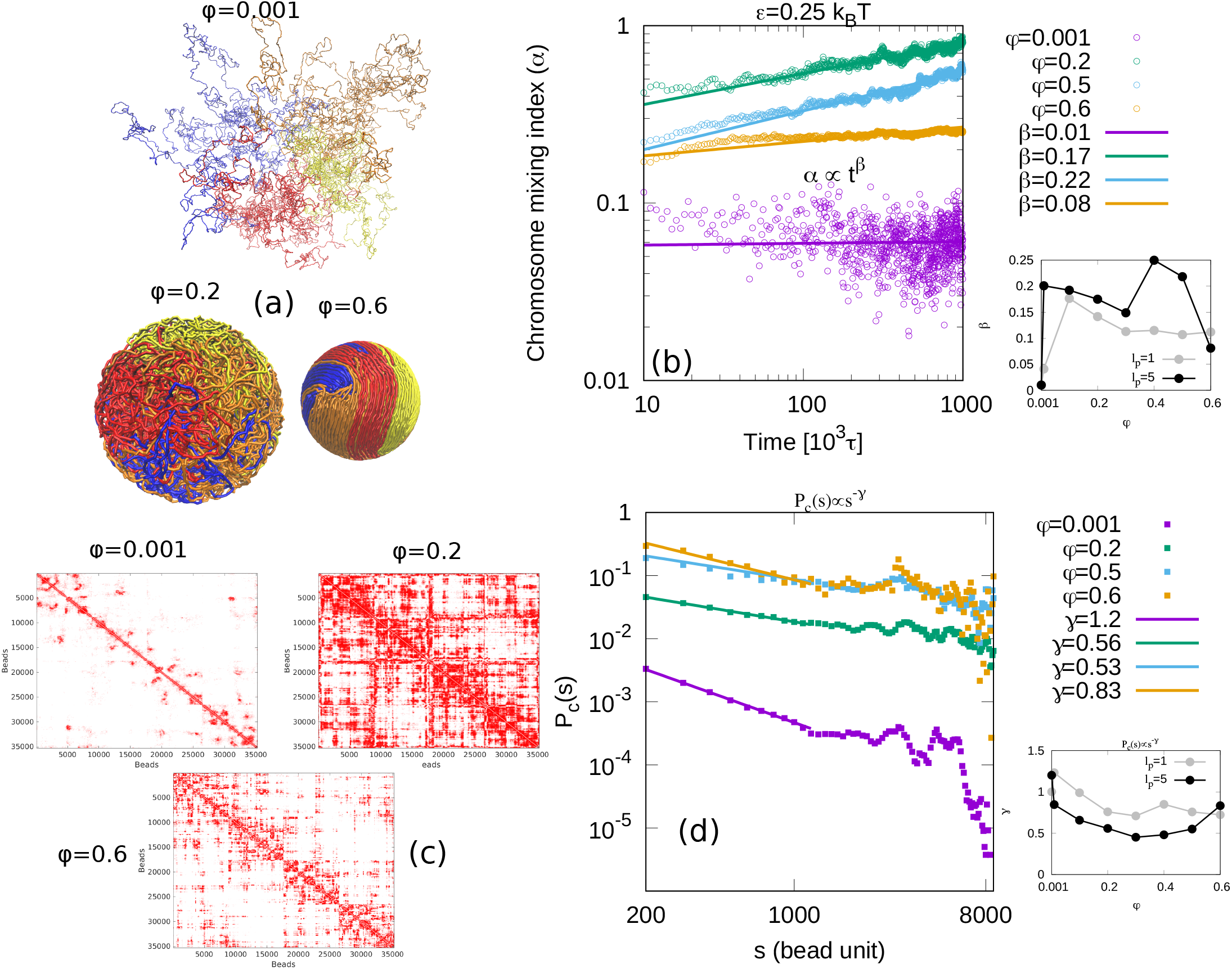
(a) Simulation snapshots are shown for different volume fractions of chromosome, *ϕ*. For *ϕ* = 0.001, corresponding to large confinement with a relatively small attraction strength (*ϵ* = 0.25), (which is below the value where collapse is observed) the chains are separated. For *ϕ* = 0.2, the chromosomes begin to mix until each chromosome approximately occupies the entire confinement volume. For *ϕ* = 0.6, the smaller volume restricts the motions of the highly condensed chains. (b) The chromosome mixing index as a function of time shows a power-law increase with the exponent *β* (*α* ∝ *t*^*β*^). Right panel of (b) The exponent first increases and then decreases as the the chain volume fraction is increased. These power laws suggests that the mixing dynamics is slow for very large and very small volume fractions (small and large confinements, respectively). (c) Contact maps obtained from the simulations, calculated by averaging the last 500 frames, are shown for the different volume fractions, *ϕ*. (d) The single-chain contact probability (*P*_*c*_(*s*)) is plotted with different colored points for different volume fractions of the chains, *ϕ*. The line corresponding to each color indicates a power-law relationship. Right panel of (d) is the scaling exponent *γ* as a function of chromosome volume fraction *ϕ* for persistence lengths *l*_*p*_ = 1 (gray color) and *l*_*p*_ = 5 (black color).

### D. Extrapolated time for full mixing of chains

We now simulated chain mixing as a function of both the volume fraction (confinement) and the attraction strength. The results of simulations for each pair of volume fractions (confinement) and chain-chain attraction strengths (*ϕ, ϵ*) are shown for *t* = 10^6^*τ* time-steps where *τ* = 60 µs; this correspond to about 60 second of real-time (see the methods section IV B). The chromosome mixing index did not reach its maximal value (*α* = 3 for 4 chains) for any pair of (*ϕ, ϵ*), which means that the chains did not completely mix during this time interval. From these simulations, we used an extrapolation of the power-laws for the mixing index as a function of time to estimate when the four chains will be completely mixed. From the relation *α* ∝ *t*^*β*^, we extrapolated the time (t) for which *α*(*t*) ≈ *α*_*max*_ = 3 where *α*_*max*_ is the ideal limit of *α* for which chains are fully mixed. In the heat map shown in Fig. 4 (a), we present the extrapolated time for the chains to fully mix for different pairs of (*ϕ, ϵ*). The heat map displays the time in seven colors according to the VIBGYOR colors pattern for the various pairs of (*ϕ, ϵ*); this shows which pairs show complete mix and on which time scale. From our calculations, we see that the pair *ϕ* = 0.4 and *ϵ* = 0.3 shows fastest mixing time *t* = 6.2 × 10^7^*τ*, which correspond to 1 hour in real time (greater than typical cell cycle time of *Drosophila* which is 8 minutes [14]). Interestingly, the value *ϵ* = 0.3 corresponds to the attraction at which the second virial coefficient of the LJ potential goes to zero (see Fig. A.3).

**FIG. 4:**
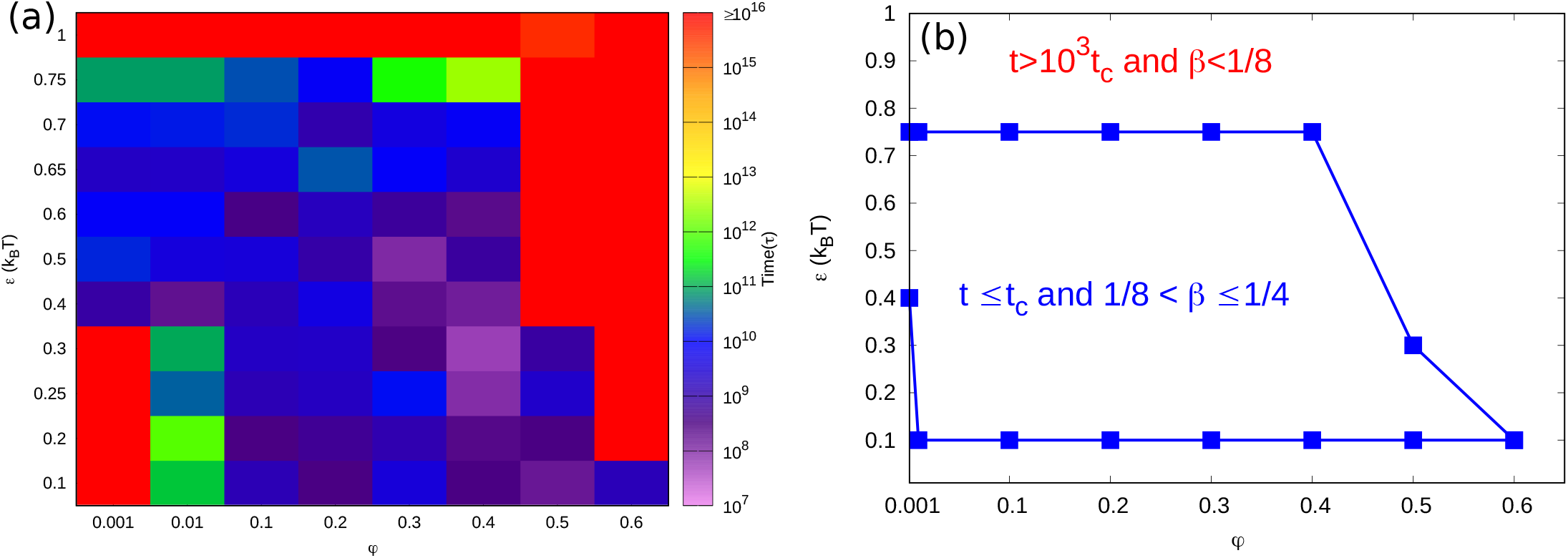
(a) Heat map of mixing times for different pairs (88 pairs) of volume fractions (confinement) and self-attraction (*ϕ, ϵ*) where 0.001 ≤ *ϕ* ≤ 0.6 and 0.1 ≤ *ϵ* ≤ 1,. The different colors indicate the order of the time when the four chains in our model are approximately maximally mixed (*α* ≤ 3). Various colors represent different times when chromosomes completely mix, and the time difference between the mixing order times that are shortest for the blue color and are longest for the red color is about 1000. Extremes of blue color region for which pair *ϕ* = 0.4 and *ϵ* = 0.75 completely mix in time *t* = 2.68 × 10^12^*τ* corresponds to 5 years and we denote it as the critical time. Note that the red color signifies extremely slow mixing, three orders of magnitude in time longer than *t*_*c*_, while the other colors show mixing on times shorter than (due to overlap with no reptation) or on the order of (due to reptation) *t*_*c*_. (b) State diagram showing the transition from slow mixing (*t* ≤ *t*_*c*_) to almost no mixing within *t*_*c*_ (*t > t*_*c*_) of the chains. In figure (b), the bars are the simulation results while the line is a guide to the eye. In the heat map (a), we observe violet, indigo and blue regions for which the chains are completely mixed by confinement and/or self-attractions for times shorter than the critical time (*t*_*c*_) and the red region for which the mixing time (due to confinement and/or self-attractions) is 3 orders of magnitude longer than the critical time.

These calculations suggest that chromosome mixing dynamics is a slow process and that most cells will probably divide before chromosomes are completely mixed. Of course, partial mixing does occur as discussed above and in [3, 18]. Different orders of magnitudes of the mixing time correspond to the different colors (from violet to red) in the heat map. The violet to green colors in heat map correspond to the range of mixing times of the order of 10^7^ to 10^12^*τ* for different pairs of (*ϕ, ϵ*). Since yellow and orange do not appear in the heat maps, this means that the chains do not mix within the times of order of 10^13^ to 10^14^*τ* for any pair of (*ϕ, ϵ*) values that were studied. In the heat map, the red color indicates the largest order of magnitude of time (*t* ≥ 10^15^*τ*) for complete mixing; for smaller times, points that are red in the heat map rarely mix. Based on the heat map colors, we can divide the time scales into two regimes: (1) slow mixing that eventually fully mixes in a long time *t*_*c*_ (to be defined), and (2) rare mixing that never fully mixes in a long time, *t*_*c*_. The time in which chains mixes within a time less than 10^12^*τ* is taken to define the critical time (*t*_*c*_ = 2.68 × 10^12^*τ* which applies to the pair *ϕ* = 0.4 and *ϵ* = 0.75). We choose this value because corresponds to 5 years and is similar to previous work where the reptation time for *Drosophila* genome is estimated as being 5 years [13]. There is a jump of 3 orders of magnitude in the mixing time for pairs of (*ϕ, ϵ*) which have longer mixing times. In Fig. 4 (b), a state diagram showing the regions for pairs of (*ϕ, ϵ*) in which chromosomes mix, albeit slowly, (*t* ≤ *t*_*c*_) and and for pairs where chromosomes mix rarely (*t >* 10^3^*t*_*c*_). The state diagram also shows scaling exponent *β* of the chromosome mixing index and for *t* ≤ *t*_*c*_, scaling exponent *β* is in the range 1/8 to 1/4 and for *t > t*_*c*_, *β* is less than 1/8. It is interesting to note that a bead of chain moves with a mean square displacement that scales *t*^1/2^ for Rouse diffusion and with mean square displacement that scales *t*^1/4^ in reptation motion [9, 10]. The root mean square (RMSD) displacements thus scale with the exponents 1/4 and 1/8 that correspond to the values obtained from the mixing index. These results suggest that the contact map dynamics (that show mixing) can be understood in terms of the single bead dynamics for the limits of non-interacting chains (Rouse dynamics with RMSD of 1/4) and strongly interacting and condensed chains (reptation dynamics with RMSD of 1/8).

### E. Effects of the lamina on the mixing of *Drosophila* chromosomes

The generic model shown above studied the mixing of 4 polymers of the same size. Here we model in a more realistic manner the genome of *Drosophila* and account for the sizes of its four chromosomes. These are denoted as Chr2, Chr3, Chr4 and ChrX, with sizes of 60.5, 68.8, 4.5 and 42.4 Mbps, respectively [53]. We modeled the *Drosophila* chromosomes as different sized bead-spring polymers (Chr2: 12,100 beads, Chr3: 13,760 beads, Chr4: 900 beads, and ChrX: 8,480 beads), with each bead corresponding to 5 kbps of DNA (see Fig. 5 (a)). In the previous sections, we have shown that open polymers (*ϵ* ≤ *ϵ*_*c*_ below the collapse transition) mix slowly in relatively small confinement volumes (*ϕ* ≥ 0.1), while strongly attractive and hence highly condensed polymers (*ϵ* = 1) rarely mix in experimentally reasonable times. In *Drosophila*, each chromosome has both relatively open (euchromatin) and relatively condensed (heterochromatin) domains (see Fig. 5 (a)). To treat this situation, we extend our previous results for homopolymers to heteropolymers that comprise blocks of both open and condensed polymers and investigate how the heterogeneity affects the mixing of several chains confinement. Each heterogeneous chromosome (blocks of euchromatin and heterochromatin) was modelled by two types of beads with different self-attractions corresponding to euchromatic and heterochromatic regions. The attraction between any two “euchromatic” beads is taken as *ϵ*_EE_ = 0.25 k_B_T and between any two “heterochromatic” beads is *ϵ*_HH_ = 1 k_B_T. In the simulation snapshot in Figs. 5 (b), euchromatic regions of the different polymers mix, while their heterochromatic regions do not. The contact map in Figs. 5 (c), calculated from the block copolymer model shows more than 4 regions along the diagonal (as in Fig. 1 (a)) due to the separate intrachain mixing of the euchromatic and heterochromatic blocks, corresponding to the chromosomes in Fig. 5 (a). From these results, it is clear that heterochromatin regions in the genome mix only rarely due to their large self-attraction; this serves to decrease the overall mixing of the chromosomes compare to homopolymers with smaller attraction strengths.

**FIG. 5:**
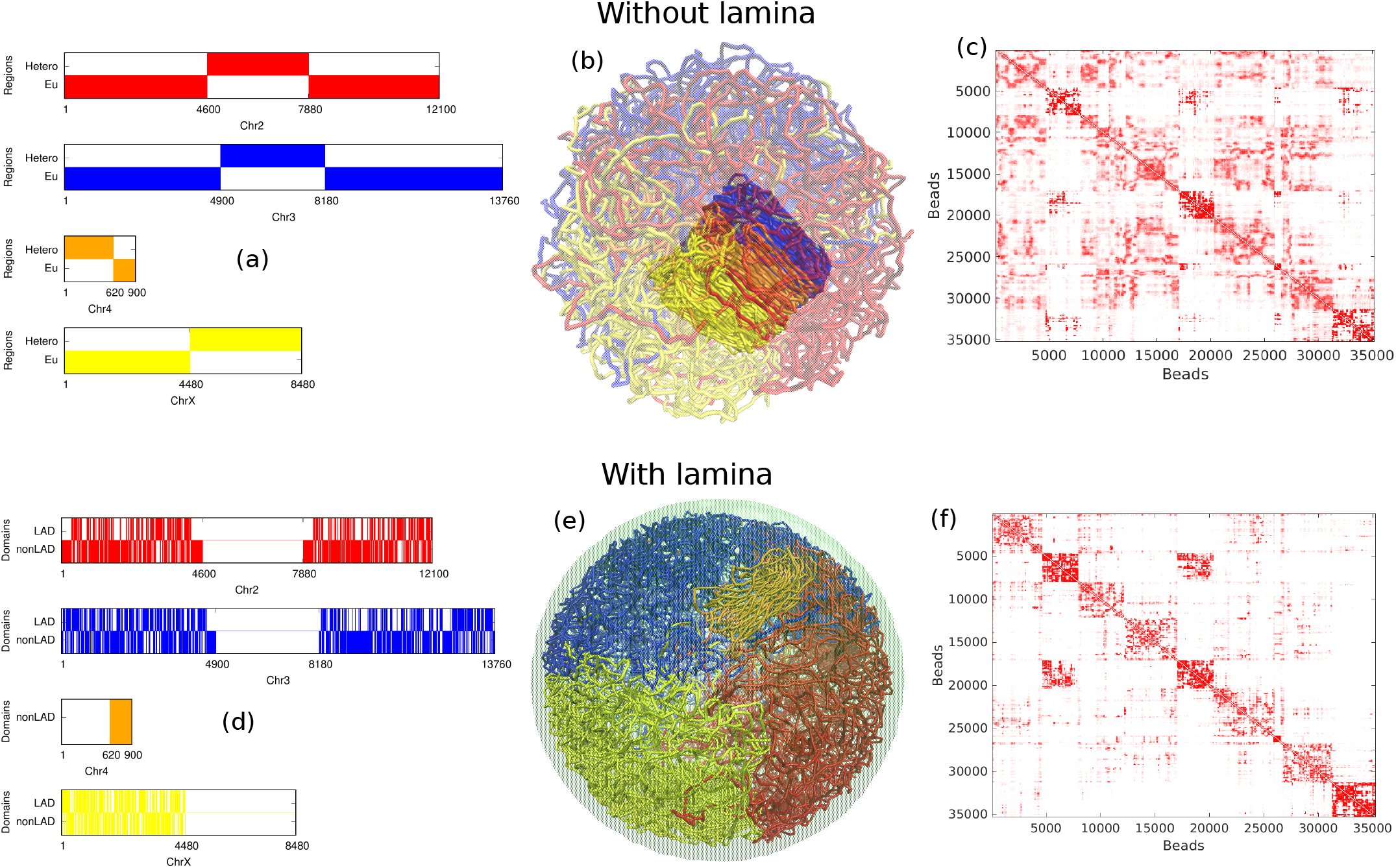
We model the chromosomes of the *Drosophila* genome by chains with two types of beads with different attraction strengths (see text), corresponding to the euchromatic (relatively small attraction) and heterochromatic (relatively strong attractions) regions. (a) Distribution of euchromatic and heterochromatic regions along each chromosome from the experimental data [53]. (b) Simulation snapshot showing chain mixing from the simulation using our block copolymer model. Euchromatic beads are transparent so that only the heterochromatic beads are visible. We observe that heterochromatic regions of the chains do not mix due to their large self-attraction. (c) The contact map calculated from the block copolymer model shows more than 4 regions along the diagonal (as in Figs. 1) due to the separate intra-chain mixing of the euchromatic and heterochromatic blocks, corresponding to the distribution shown in (a). To understand the role of the binding of the chromatin to the lamina in chromosome mixing, we modeled the bonding of certain chromosome segments (LADs) to the nuclear lamina using our block copolymer model for *Drosophila*. (d) The distribution of LAD and non-LAD domains along each chromosome based on the experimental data[54]. (e) Snapshot of simulation for chromosome volume fraction *ϕ* = 0.1, in which in presence of lamin (green), the four chromosomes Chr2 (red), Chr3 (blue), Chr4 (orange) and ChrX (yellow) of *Drosophila* genome separate into distinct regions and do not mix. (f) The contact maps for case (e). Comparing this to the contact map for the same block copolymer model in the absence of binding to the lamina in Fig. 5 (c), we observe a smaller probability of contact in the off-diagonal regions. From these results, it is clear that the binding to the nuclear lamina decreases mixing of chromosome.

Experimental studies have shown that specific chromosome regions, denoted as lamin associated domains (LADs), distributed along the chromosome [45] are tethered to the nuclear lamina. To account for the role of lamina in chromosome mixing, we included the bonding of some fraction of the beads (representing the LADs) to the nuclear lamina within our block copolymer model for *Drosophila*. The nuclear lamina is modeled as a thin layer of “lamin beads” localized near the surface of the spherical confinement. In Fig 5 (d), we have shown the distribution of LAD (lamin associated domains) and non-LAD domains along each chromosome based on the experimental data [54]. Note that this data is available only for the euchromatic regions of the chromosomes. Besides LADs and non-LADs, each chromosomal bead corresponds to either euchromatin or heterochromatin according to our block copolymer model. Figs 5 (e) shows a simulation snapshot for chromosome volume fraction *ϕ* = 0.1. We see that the chains separate in distinct territories and rarely mix due to lamin-LAD attraction at the nuclear periphery. We have calculated the contact map in presence of lamin and compared it to the contact map for the same block copolymer model in the absence of binding to the lamina in Fig 5 (f). In the presence of the lamin binding, the simulations show less contact in the off-diagonal regions. From these results, it is clear that the binding to the nuclear lamina reduces the mixing of chromosome.

## III. DISCUSSION

Our findings suggest that the chromosome mixing index *α*(*t*) is useful as a single, time dependent parameter that quantifies the slow (and sometimes, extremely slow) mixing of different chromosomes confined to the nucleus. For ideal chains within small confinement, the mixing index of chromosomes is maximal; a given chromosome mixes with the *n* − 1 other chromosomes. The value of chromosome mixing index depends on the volume fraction and chromosome interactions that change the scaling law of how chains mix as a function of time. The chromosome mixing index increases slowly over time: *α* ~ *t*^*β*^ with an exponent of *β*. For slowly mixing chromosomes, our simulations predict a mixing exponent of *β* = 1/8 − 1/4 depending on (*ϕ, ϵ*). Unconfined polymers collapse when their self-attraction strength is less than a critical strength (*ϵ < ϵ*_*c*_) related to the vanishing of the second virial coefficient at the theta-point at which the polymer first collapses (see Fig. A.3). In contrast, confined polymers collapse when their confinement radius is smaller than the radius of gyration of the unconfined polymer (*R*_*c*_ " *R*_*g*_).

We used the same dynamical simulations to calculate the scaling exponent of contact probability, *P*_*c*_(*s*), for segments of single chain separated by a contour length *s*: *P*_*c*_(*s*) ~ *s*^*−γ*^ with *γ* ≤ 1 for collapse chains, independent of whether they collapse due to attraction strength (*ϵ*) or due to small confinement (relatively large volume fraction, *ϕ*). Open, weakly or non-attractive polymers (as a model of euchromatin) mix even in relatively small confinement (nucleus), while strongly attractive polymers that collapse and become highly condensed (as a model of heterochromatin) mix extremely slowly. Hi-C experiment generate, contact maps of both single chromosomes as well as the entire genome. The Hi-C contact map of a single chromosome is helpful in identifying the sequences that result in A/B compartments, TADs, and loops within a chromosome. In contrast, the Hi-C contact map of the entire genome is usually used to identify the distinct chromosome territories; its off-diagonal contacts are often overlooked because these typically show less contact than the diagonal elements. However, it is precisely these off-diagonal contacts that are indicative of the extent of chromosome mixing. We therefore defined and have focused on the chromosome mixing index which is the ratio of the off-diagonal contact to diagonal contacts and thereby provides information on the degree of chromosome mixing. Recent studies have indeed observed the overlap of chromosome territories, suggesting that the different chromosomes are mixed together at their borders territories [3–7]. This has important implications for the correlation of gene activity of different chromosomes [8].

The overlapping of partially mixed chains (which represent chromosomes) can be seen in our simulation results by focusing on two chains Chr2 and Chr3 from the case Fig. 6 (a) and counting the number of beads of Chr2 the lie within the overall contour of Chr3 and vice versa (see Fig. 6 (b)). The overlap of these chains at their edges indicates partial mixing. Fig. 6 (e), we have calculated the chromosome mixing index from the Hi-C experiment data of *Drosophila* genome [16]. The chromosome mixing index we calculate from the experimental data *α* = 0.3, and its maximum value was calculated theoretically from a generalization of equation in the methods section IV E (to chromosomes of different sizes) as considering heteropolymers, which is approximately 4 (*α*_*max*_ = 3.9). Comparing this with our simulation results for the case of Fig. 6 (a), we find that the simulations predict a chromosome mixing index of *α* = 0.24 (see Fig. 6 (d)). This indicates that extension of the theoretical model to include different euchromatin/heterochromatin interactions and LAD/non-LAD interaction with the lamina yields mixing whose index is comparable to Hi-C experiment data. The scaling exponent of the contact probability factor *γ* is useful to understand the mixing of subdomains within single chains, whereas the scaling exponent of the chromosome mixing index *β* is useful to understand the mixing of different chains.

**FIG. 6:**
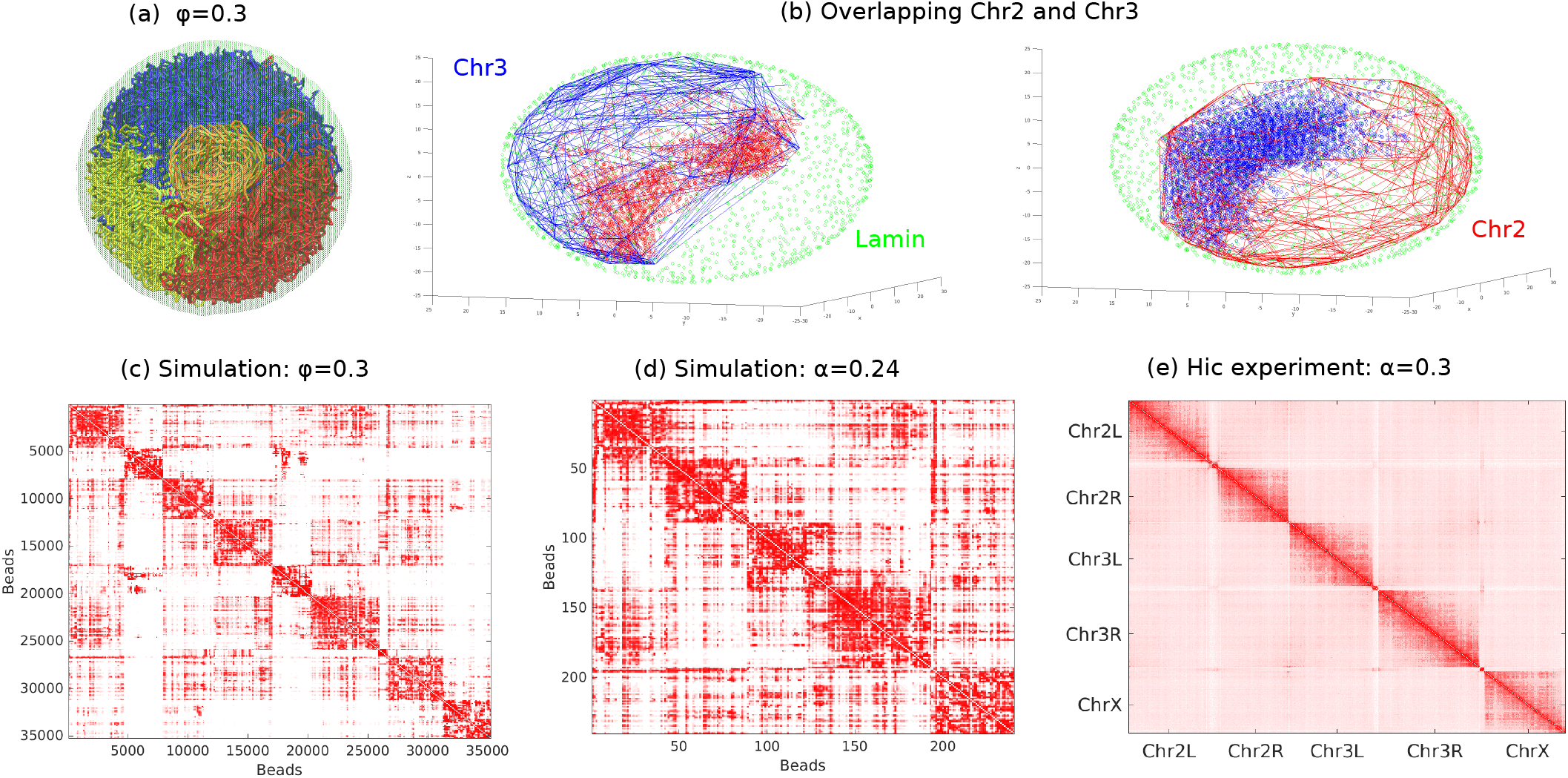
(a) Snapshot of simulation of block copolymer model in presence of nuclear boundary (lamin) attractions for a chain volume fraction of *ϕ* = 0.3 (b) Overlapping between Chr2 and Chr3 are shown. Lamin beads are shown by green points. Chr2 and Chr3 regions in 3D are shown by the line and overlapping regions of Chr2 and Chr3 are shown by red and blue points. (c) The contact map for case (a). (d) The contact map and chromosome mix index are calculated for the simulations (a) by omitting the heterochromatin regions. (e) Contact map and chromosome mixing index are calculated from the Hi-C experiments data of *Drosophila* genome [16]. Note that *Drosophila* Hi-C experiments contact map data is only available for euchromatic regions.

We note that it is possible that there are only a few universality classes for these scaling exponents. For example, the dynamical exponent may have one value when Rouse dynamics is applicable (root mean squared displacement (RMSD) of a bead varies as *t*^1/4^ and another for the strongly attractive regime when the chains are highly condensed and the dynamics occur via reptation (RMSD varies as *t*^1/8^). Indeed these are the values for the dynamical mixing exponent *β* the for theta-point (vanishing of the second virial coefficient) and strongly attractive chains respectively (see Table IV H for comparison between scaling exponent of RMSD and scaling exponent of mixing index). This agreement indicates how the polymer dynamics via the RMSD exponents for bead motion relates to the dynamical evolution of the entire contact map – an important link between the polymer physics and the experimental Hi-C biology. The variation of the exponent between these limits may be due to crossover effects, although we note predictions of the RMSD of interacting polymers that are neither Rouse-like nor collapsed give scaling exponents that vary with the classes of interactions [52].

## IV. METHODS

### A. Generic Model

To study the mixing of chromosomes in *Drosophila*, we simulate the confinement of 4 chains of equal length, where each chromosome is modeled as coarse-grained bead-spring polymer chain with *M* = 8810 beads; this gives a total *N* = 35240 beads in all 4 chains. Each bead represents 5 kbps of DNA and the associates histones, corresponding to a bead diameter of *σ* = 30 nm (see the methods section IV B). The beads of each chain are connected to their nearest neighbors along the contour of the chain, by “springs” (harmonic potential) and the energy is written:

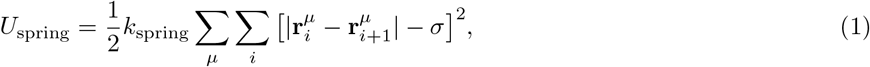

where 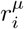 is the position of *i*^th^ bead of the *µ*^th^ chain with *µ* = 1… 4. *k*_spring_ is the spring constant. Each chain has a persistence length of 5 beads, based on a previous estimate that interphase chromosomes consist of compact fibers with a diameter which is approximately 30 nm with a persistence length of 150 nm [55]. We also simulated chains with a persistence length of 1 bead and compared the results with those whose persistence length was 5 beads. Since the bending energy determines the persistence length [56], this estimate allows us to write the bending energy of chromosome as:

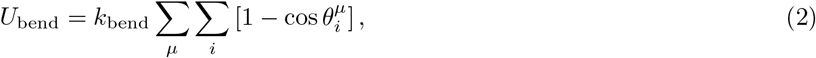

where *θ*_*i*_ is the angle between three adjacent beads within the chain. *k*_bend_ is the bending stiffness of the chain which is related to the persistence length by 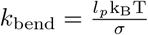 [56]. Here *l*_*p*_ is the persistence length, *k*_B_ is Boltzmann constant, and *T* is absolute temperature. Apart from the nearest-neighbour spring interaction described above, any two beads (that can be anywhere along the contour of the chain) interact via the standard Lennard-Jones (LJ) potential (*U*_LJ_) that depends on their three-dimensional spatial separation, with an energy:

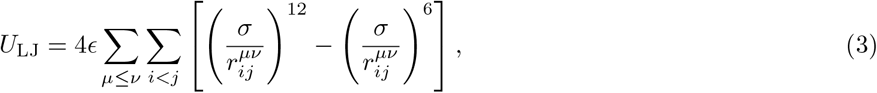

where 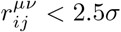 and zero otherwise. Here, 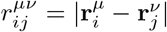 is the distance in 3D space between *i*^th^ and *j*^th^ beads of any chain and *ϵ* is strength of potential. We varied *ϵ* from 0 to 1 which can then account for no interactions (*ϵ* = 0 for phantom chains), repulsive interactions (small values of *ϵ*), and attractive interactions (larger values of *ϵ*) between any two beads. The chains are confined by a impenetrable, spherical wall of radius *R*_*c*_ that mimics the effect of the nuclear envelope. A confinement potential *U*_nucleus_ is used to account for the hard-core repulsion between the beads and the spherical wall. We define the parameter *ϕ*, as the volume fraction of chains within the sphere (corresponding to chromosomes in the nucleus)

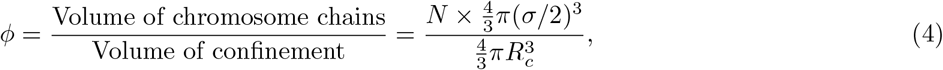

*ϕ* ranges from 0.001 to 0.6, where small values of *ϕ* represent chains in large confinement and large values represent chains in small confinement. We performed simulations for *ϕ* = 0.001, 0.01, 0.1, 0.2, 0.3, 0.4, 0.5 and 0.6 which are respectively equivalent to confinement radii of *R*_*c*_ = 164, 77, 36, 29, 25, 23, 21 and 20 in bead units (*σ*). The total energy of the system is given by the following equation:

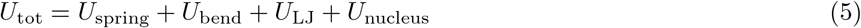

To simulate the system, we used the molecular dynamics simulation package LAMMPS [57], in which Newton’s equations are solved for particles in a solvent, represented by a motion in the presence of a viscous force and a Langevin thermostat to ensure an NVT ensemble. To obtain generic results, we first simulate chromosomal chains using the polymer model described above. Later on, we modified the model to more realistically model the *Drosophila* genome. To do this, we included heterogeneity within each chain to represent the block copolymer structure of a chromosomes with both euchromatin/heterochromatin, the presence of LAD /non- LAD chromosome domains, and attractive interactions between the LAD domains and the nuclear lamina.

### B. Physical units of chromosome mixing models

In our coarse-grained model, each chromosomal bead represents 5 kbps of DNA, which represents 25 nucleosomes and their linker DNA. A nucleosome is modeled as cylindrical structure with a diameter of 11 nm and a length of 5.5 nm [58]. The volume of 25 nucleosomes is comparable to the volume of a spherical bead of diameter to be 30 nm; therefore, we choose the diameter of chromosomal bead *σ* = 30 nm. The time it takes such a bead to diffuse its own size (*σ*), is denoted as the Brownian time *τ*, defined by 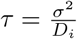, where *D*_*i*_ = k_B_ T*/ζ*, is the diffusion coefficient of *i*^th^ bead. The friction coefficient *ζ* is calculated by using the Stokes law for a spherical bead of diameter *σ* in a solution of viscosity *η* and given by *ζ* = 3*πησ*. When a bead’s diameter *σ* is known, the Brownian time is as *τ* = 3*πησ*^3^/k_B_T. Assuming that the viscosity of the the nucleoplasm is similar to water, we find that with a viscosity *η* = 1 cP [59], temperature *T* = 310*K*, and a bead diameter of *σ* = 30 nm, the Brownian time of a coarse-grained bead is *τ* = 60 µs.

### C. Initial conditions

During cell division, each chromosome condenses and separates from each other, and after cell division, two daughter cells are formed. This motivates our initial condition for which chains are initially condensed and separated in our simulation.

### D. Simulation parameters

### E. Chromosome mixing index

We define the chromosome mixing index as the ratio between the total number of interchromosomal contacts and total number of intrachromosomal contacts. This is easily calculated from the experimental Hi-C contact map. The maximum value of the chromosome mixing index is theoretically given by

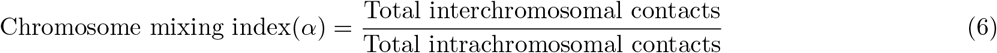

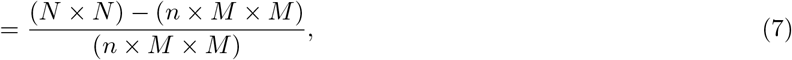

where *N* is the total size of the genome, *M* is the length of the individual chromosome, and *n* is the total number of chromosomes in the genome.

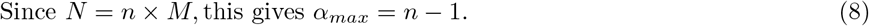

Since *N* = *n* × *M*, this gives *α*_*max*_ = *n* − 1. (8) In the ideal limit (where there are no interactions) a given chromosome can mix with any of the *n* − 1 chromosomes.

### 5. Calculating the contact map, contact probability, and chromosome mixing index from simulation data

In every 10^3^*τ* time steps (corresponding to one frame, total 1000 frames), the simulation uses the Langevin equations of motion to generate the position of each bead in Cartesian coordinates. From the Cartesian coordinates, we calculate the distance (*r*_*ij*_) between any two beads *i* and *j*. The first step in generating a contact map from the simulation data is to define the contact criteria based on the distance (*r*_*ij*_) between any two beads *i* and *j*. We choose to do this by saying that when the distance between any two beads *i* and *j* is less than 1.5 times of their minimal distance due to hard sphere repulsion (*σ*), they are considered to be in contact (*r*_*ij*_ *<* 1.5*σ*). By calculating which pairs of beads are in contact, we generate a contact matrix with *N* × *N* elements, that includes all the contact information of the *N* beads in our system. Since we do not need the details of every contact of every bead for our analysis, we coarse-grain the matrix into blocks of 500 kbps resolution, which reduces the size of the contact matrix by a factor of 100. We use the block average method to coarse-grain the contact matrix [60]. In this method, at the *m*^th^ level of coarse-graining, an *N* × *N* matrix is divided into blocks of size *m* × *m* each. Then a new contact matrix of size *N*_*m*_ × *N*_*m*_ where *N*_*m*_ = *N/m* is constructed in such a way that the value of each element represents the arithmetic mean of the elements in each block. By setting *m* = 100, we coarse-grain the original contact matrix into a 352 × 352 contact matrix. Because one contact matrix is generated per frame, 1000 frames result in 1000 contact matrices. We denote by 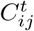 the contact matrix of the *t*^th^ frame; then, a mean contact matrix *C*_*ij*_ is calculated by averaging the contact matrices over the last 500 frames. We represent the resulting contact map (CM) as a heat map of the mean contact matrix (CM = log_2_[*C*_*ij*_]) in order to visualize the contact matrix.

We use the contact matrix (*C*_*ij*_) to calculate the contact probability (*P* (*s* =| *i* − *j* |)) between pairs of beads (*i, j*). We use the full contact matrix (*C*_*ij*_) to cut out 4 matrices (*IC*_*µ*_) along the diagonal, each of size *N*_*m*_/4, representing the contact matrix of the *µ*^th^ chromosome. We write *IC* = (1/4) ∑_*µ*_ *IC*_*µ*_ which represents the mean intra-chromosome contact matrix of all four chromosomes. The contact probability (*P*_*µ*_(*s*)) within each chain is calculated by using following formula which is then averaged over all the chains given by

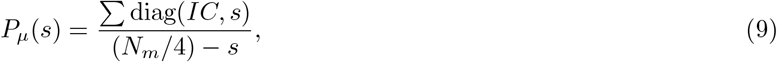

where diag (*IC, s*) are the elements of the *s*^th^ diagonal of matrix *IC*. Note that *s* = 0 represents the main diagonal and *s >* 0 is above the main diagonal.

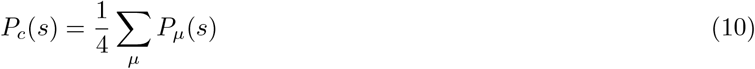

The chromosome mixing index (*α*) is calculated from the simulation as the ratio between the sum of the number of contacts between chains (**not** including the two nearest-neighbor beads to which each bead is bonded) and the sum of intra-chain contacts. The chromosome mixing index (*α*) for *t*^th^ time frame is calculated by

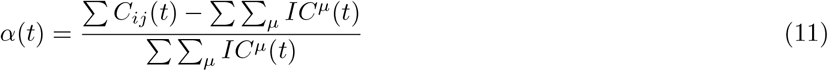

where ∑ *C*_*ij*_ is the sum of all elements in matrix *C*_*ij*_ and *t* is the time. Here, *C*_*ij*_ is the averaged contact matrix of all the chromosomes and *IC*^*µ*^ is the intra-chromosome contact matrix of *µ*^th^ chromosome, as defined above.

### G. Variation of scaling exponent of MSD

**TABLE II:**
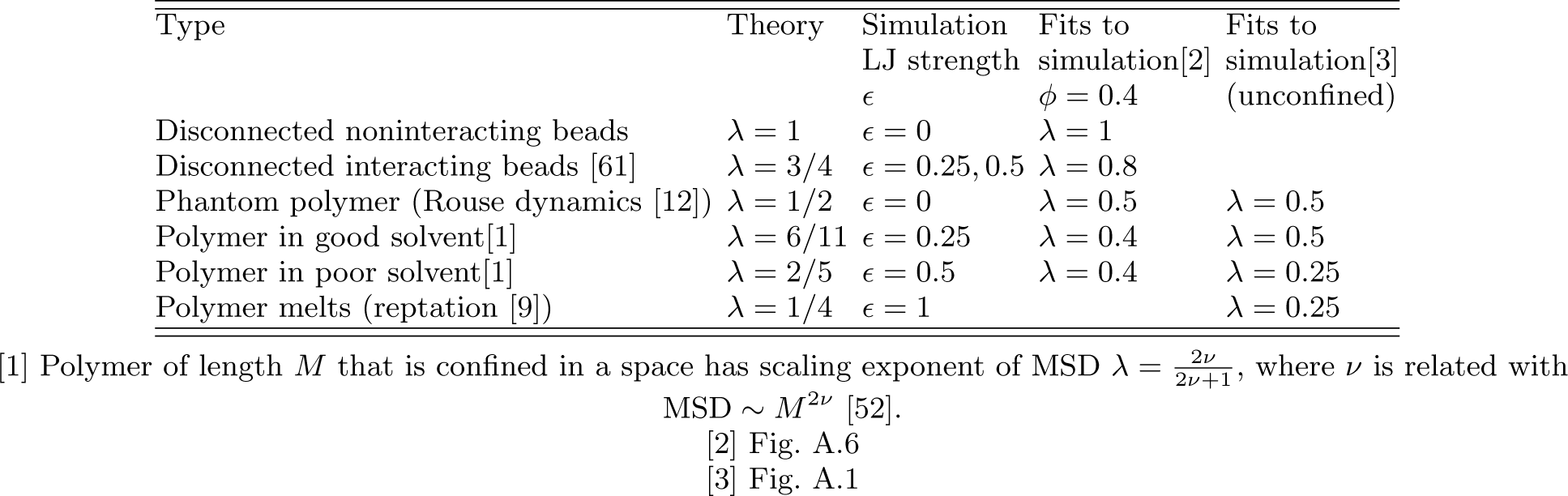
MSD, (*r*^2^) ~ *τ*^*λ*^

### H. Comparison between scaling exponent of RMSD and scaling exponent of mixing index

**TABLE III:** RMSD 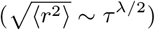 and chromosome mixing index (*α* ~ *t*^*β*^)

| Type | LJ strength<br>$\epsilon$ | $\lambda/2$<br>(for $\phi = 0.4$) | $\lambda/2$<br>(unconfined) | $\beta$<br>(for $\phi = 0.4$) |
| --- | --- | --- | --- | --- |
| Polymer in good solvent | $\epsilon = 0.25 \text{ k}_B\text{T}$ | $\lambda/2 = 0.2$ | $\lambda/2 = 0.25$ | $\beta = 0.25$ |
| Polymer in poor solvent | $\epsilon = 0.5 \text{ k}_B\text{T}$ | $\lambda/2 = 0.2$ | $\lambda/2 = 0.125$ | $\beta = 0.22$ |
| Polymer melts (reptation) | $\epsilon = 0.75 \text{ k}_B\text{T}$ | | $\lambda/2 = 0.125$ | $\beta = 0.13$ |
| Polymer melts (reptation) | $\epsilon = 1 \text{ k}_B\text{T}$ | | $\lambda/2 = 0.125$ | $\beta = 0.12$ |

## Acknowledgments

We thank Omar Adame-Arana, Dan Deviri, Amit Kumar, Shensheng Chen and Prof. Zhen-Gang Wang for useful discussions.

## Appendix A

Unconfined single chain as a function of self-attraction strength

Here, we discuss how the attraction of the beads characterized by the parameter *ϵ* affects the radius of gyration (*R*_*g*_), mean-square displacement (MSD) as a function of time, and the contact probability scaling exponent (*γ*) of an unconfined single chain. In Fig. A.1 (a), we have shown 3D snapshots of our simulations of unconfined, single chains of *M* = 8810 beads by varying the attraction strength for the cases of persistence lengths of both 1 and 5 beads persistence. To better understand how the different chain segments mix in 3D, we divide the chain into 10 different colors along its length, from blue at one end to red at the other. In Fig. A.1(b), we plot the radius of gyration of a chain with a persistence length of 1 bead as a function of the interaction strength. For *ϵ* = 0, the radius of gyration obeys the power law scaling as a function of the chain length, *M* : *R*_*g*_ ~ *M*^*?*^ with *ν* = 1/2, consistent with the pure random walk of a phantom polymer. For small values of *ϵ >* 0 to *ϵ* ≤ 0.25 the beads repel each other and *R*_*g*_ increases with *ϵ* indicative of chain swelling due to exclude volume repulsion. For *ϵ >* 0.25, *R*_*g*_ decreases with the interaction strength signifying collapse; we observe a sharp drop starting at *ϵ* = 0.25 and we estimate that at *ϵ* = 0.3 the chains undergo a collapse transition. This is consistent with an estimate of the theta-point (where the second virial coefficient [51] for the LJ potential vanishes) based on the LJ potential (see Fig. A.3). This defines the critical value *ϵ*_*c*_ of the collapse transition.

For larger values of *ϵ* (*ϵ* ≥ 0.5), the radius of gyration as a function of chain length follows the scaling exponent *ν* = 1/3 indicates that the polymer is collapsed since it is in a poor solvent. The case of the persistence length *l*_*p*_ of 5 beads is similar. Fig. A.1(c), shows that the radius of gyration follows power law scaling *R*_*g*_ ~ *l*_*p*_ (*M/l*_*p*_)^*ν*^ with *ν* = 1/2 for *ϵ* = 0 and *ν* = 1/3 for *ϵ* ≥ 0.5. for the case of the 5 bead persistence length we observed the first collapse at *ϵ*_*c*_ = 0.4 instead of *ϵ*_*c*_ = 0.3 for the case where *l*_*p*_ = 1 bead. This means a chain with *l*_*p*_ = 5 beads requires is stiffer over a longer distance and requires a larger attraction among the beads in order to collapse.

From these simulations we have calculated the contact probability that follows a scaling law *P* (*s*) = *s*^*−γ*^ and present the results of the scaling exponent *γ* as a function of *ϵ* in Fig. A.1(d). For a unconfined single chain of persistence of 1 and 5 beads, the contact probability scaling exponent is *γ* = 1.5 for *ϵ* = 0 (random walk chain). This is indeed consistent with the behavior of a completely random, Gaussian chain where the normalization of the probability yields *γ* = 3/2. In Fig. A.1(d), we plot the contact probability scaling exponent *γ >* 1 for an open chain (*ϵ < ϵ*_*c*_) and *γ* ≤ 1 for a collapsed chain (*ϵ* ≥ *ϵ*_*c*_). For the collapsed chain the values of 0 *< γ <* 1 signify intermediate level mixing while *γ* = 1 shows no mixing of different parts of chain that are far apart.

In addition, we also calculated the mean-square displacement (MSD) of a bead (averaged over all the beads of the chain) by varying the LJ attraction strength (see Figs. A.1(e),(f)). For *ϵ* = 0 and 0.25 when chain is open, MSD ~ *τ* ^1/2^ and bead moves like a monomer in a Rouse chain [12]. For *ϵ* = 0.5, 0.75, 1 when chain is strongly collapsed, MSD ~ *τ* ^1/4^ and bead moves like a monomer in a reptating chain [9]. Every bead in a collapsed chain must cross the crowded environment of other beads in order to move, so the reptation model is more appropriate. We have also shown these results do not change by changing chain’s persistence length from *l*_*p*_ = 1 bead to *l*_*p*_ = 5 beads (see Figs. A.1(e),(f)).

**FIG. A.1:**
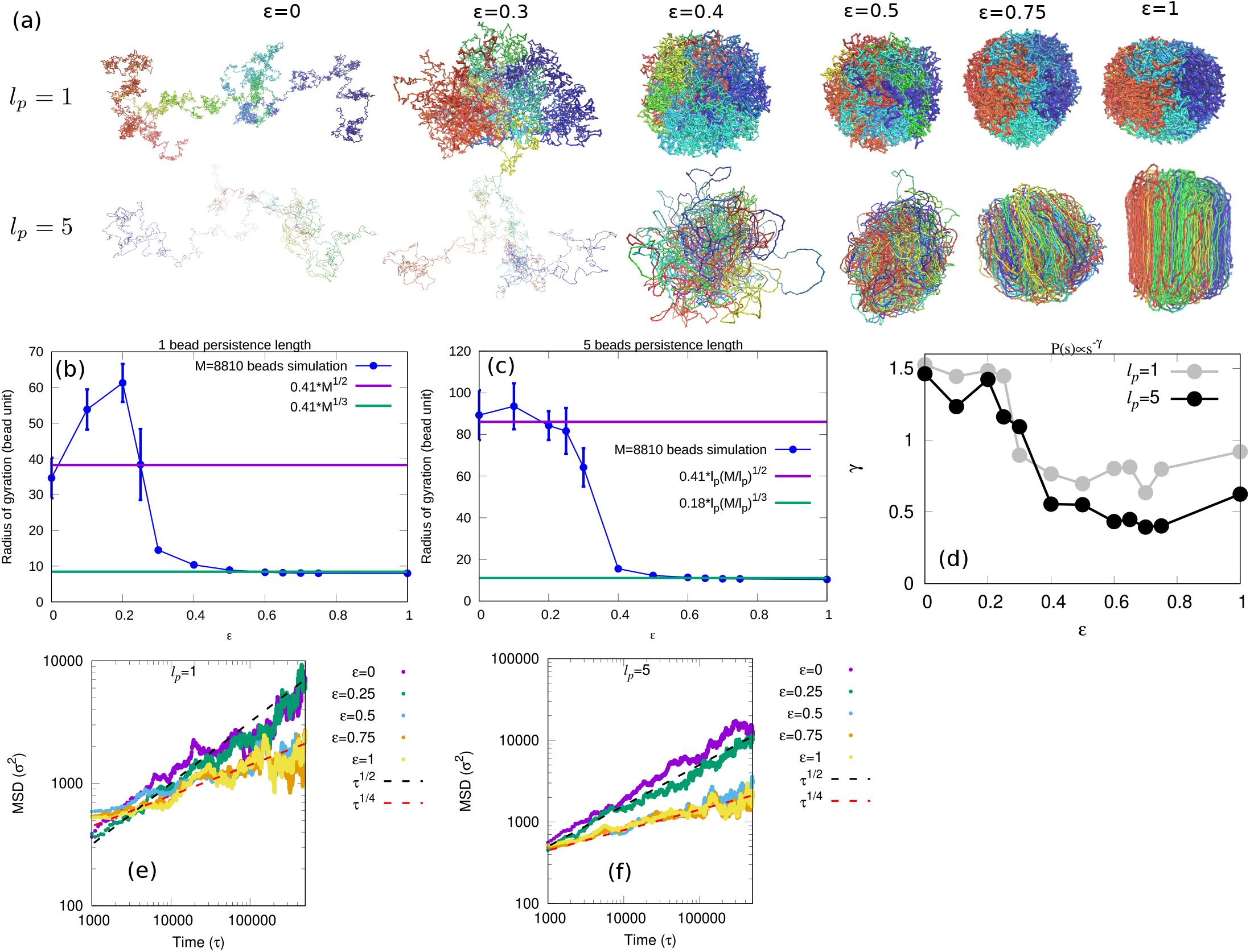
(a) Late time snapshots (*t* = 10^6^*τ* time steps) of simulated, unconfined single chains for persistence lengths of *l*_*p*_ = 1 and *l*_*p*_ = 5 beads as a function of the LJ interaction strength, *ϵ*. The chain is colored from blue to red along its length to exhibit the mixing of different segments of the chain. Radius of gyration as a function of LJ attraction strength for the persistence length (b) *l*_*p*_ = 1 beads and (c) *l*_*p*_ = 5 beads. Simulated results are compared with scaling law of radius of gyration (*R*_*g*_ ~ *M*^*ν*^). (d) Contact probability scaling exponent as a function of *ϵ* for *l*_*p*_ = 1 bead (gray color) and *l*_*p*_ = 5 bead (black color). Mean-square displacement (MSD) of a bead (averaged over all the beads of the chain) is calculated for the persistence length (e) *l*_*p*_ = 1 bead and (f) *l*_*p*_ = 5 beads. In both figures (e) and (f), the MSD fit to *τ* ^1/2^ (black dotted line) for *ϵ* = 0 and 0.25 and fit to *τ* ^1/4^ (red dotted line) for *ϵ* = 0.5, 0.75, 1.

**FIG. A.2:**
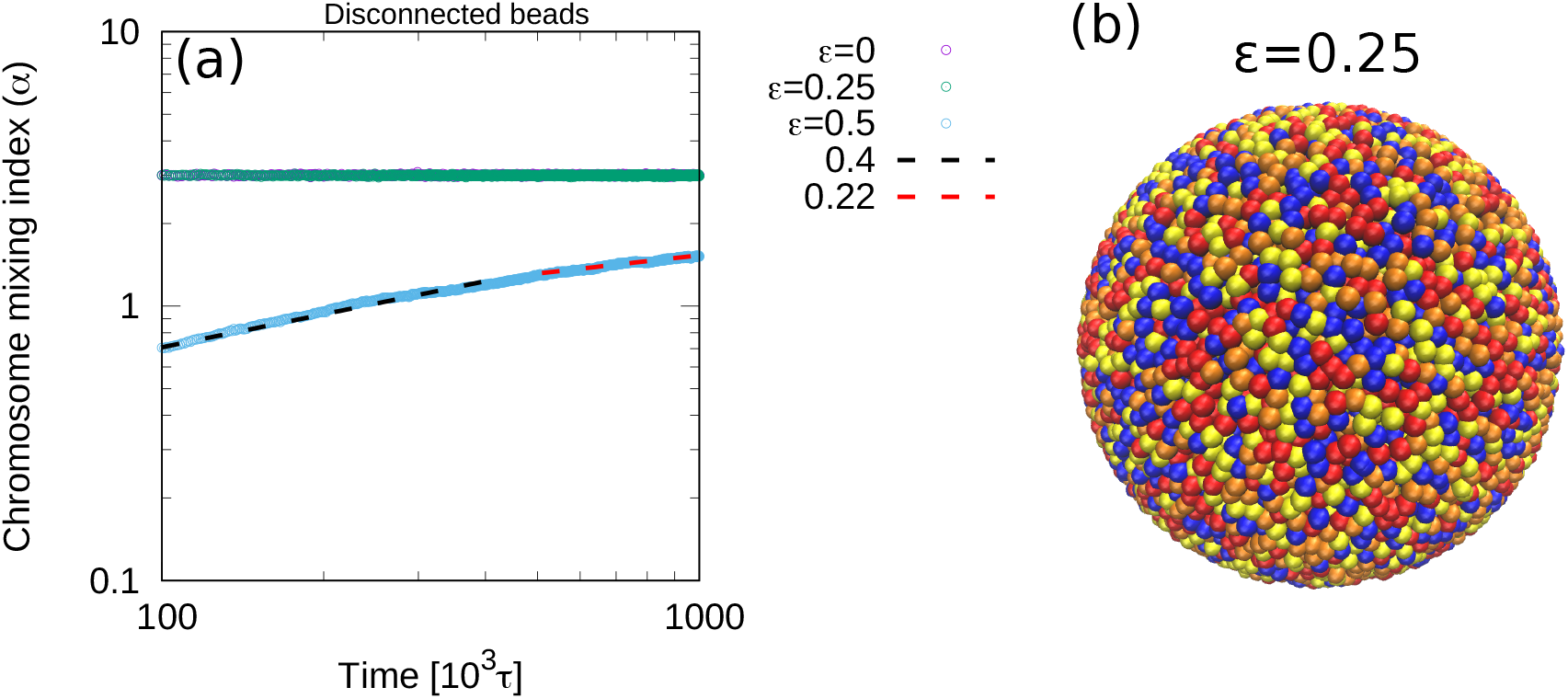
Mixing of disconnected chromosome beads: Simulation results are shown for volume fraction *ϕ* = 0.4. (a) Mixing index *α* are shown for different *ϵ*. For *ϵ* = 0.25, disconnected beads mix quickly and reach the maximum value of mixing index (*α* = 3). (b) Late time snapshot (*t* = 10^6^*τ* time steps) of simulated disconnected chromosome beads for attraction strength *ϵ* = 0.25.

**FIG. A.3:**
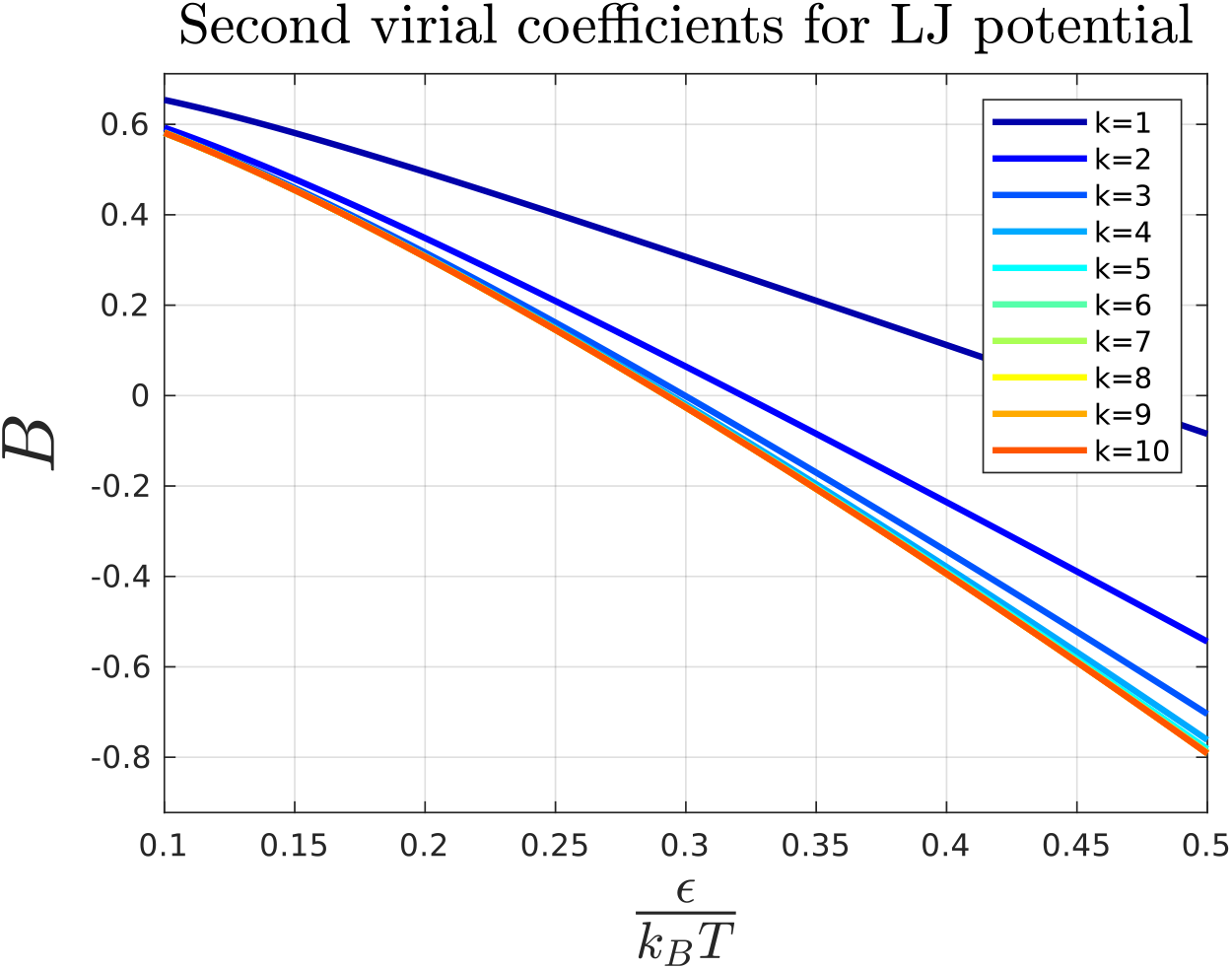
Second virial coefficient (SVC) of the Lennard-Jones potential: The LJ potential describes both the attraction and repulsion between non-bonded beads. SVC is used to determine the value of *ϵ* for which repulsive and attractive bead-bead interactions are equal (theta solvent condition). The integral for *B*(*T*) cannot be calculated analytically, but an accurate approximation can be derived from the 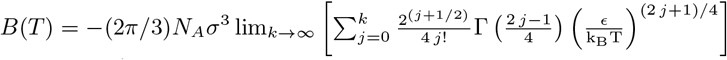[51] In the figure, the function *B*(*T*) is plotted as a function of *ϵ /*k_B_T for *k* = 1 to 10. For larger value of k, the graph saturates and at *B* = 0, we get *ϵ*_*θ*_ = 0.2925 k_B_T.

**FIG. A.4:**
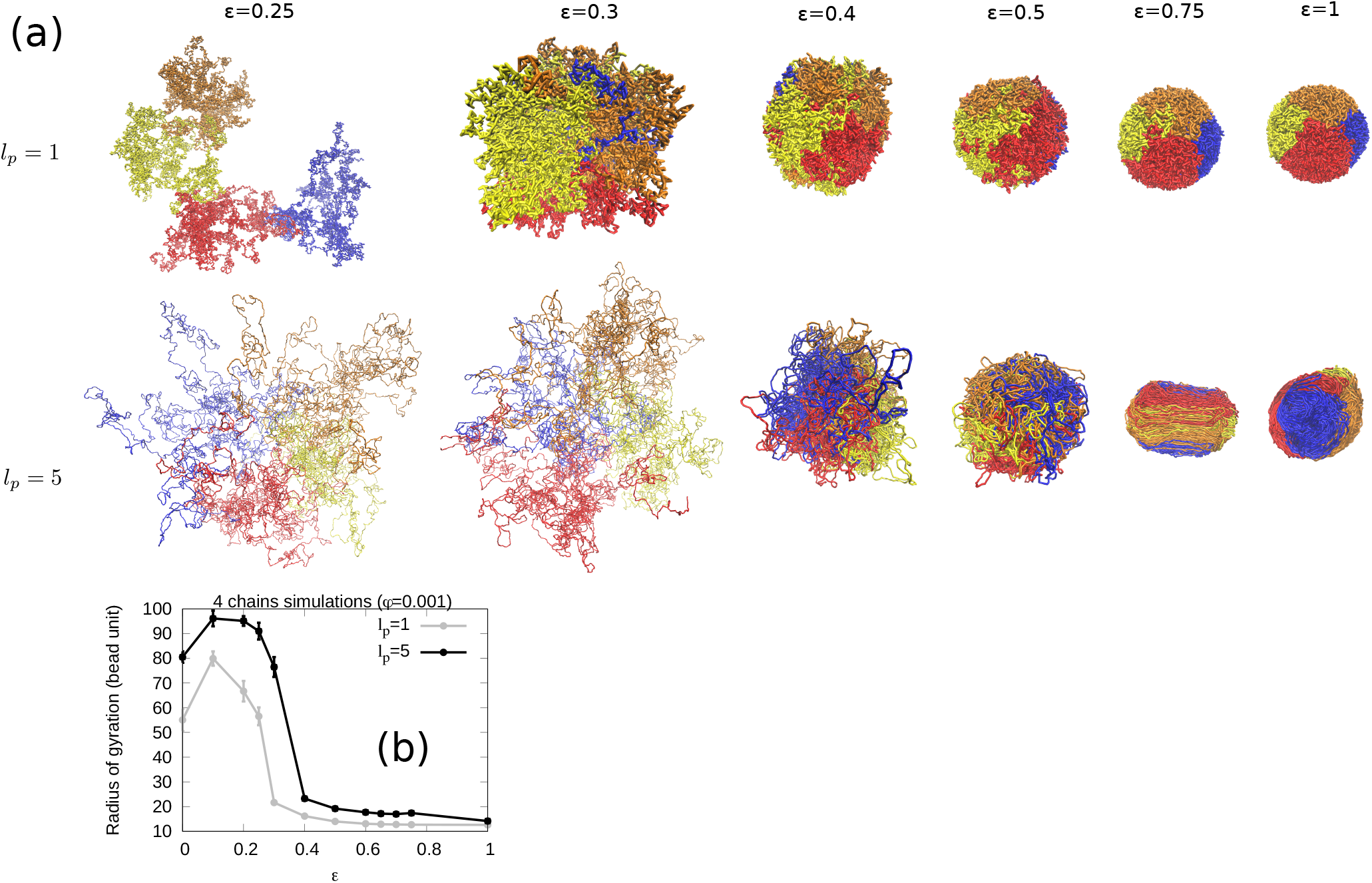
Mixing of four chains in large confinement volume (corresponding to a small volume fraction of beads, *ϕ* = 0.001) while varying the bead-bead interactions, *ϵ* : (a) Simulation snapshots showing the mixing of four chains mixing as a function of the interaction strength, *ϵ*, for persistence lengths *l*_*p*_ = 1 bead and *l*_*p*_ = 5 beads. Note that some snapshots were zoomed out because they were too large and would take up too much space if they were shown at their actual size. (b) Radius of gyration as a function of *ϵ* for *l*_*p*_ = 1 bead (gray color) and *l*_*p*_ = 5 beads (black color). Note that attraction strength at which collapse occurs for the one-bead persistence length is *ϵ*_*c*_ = 0.3 whereas for the 5-beads persistence length, *ϵ*_*c*_ = 0.4.

**FIG. A.5:**
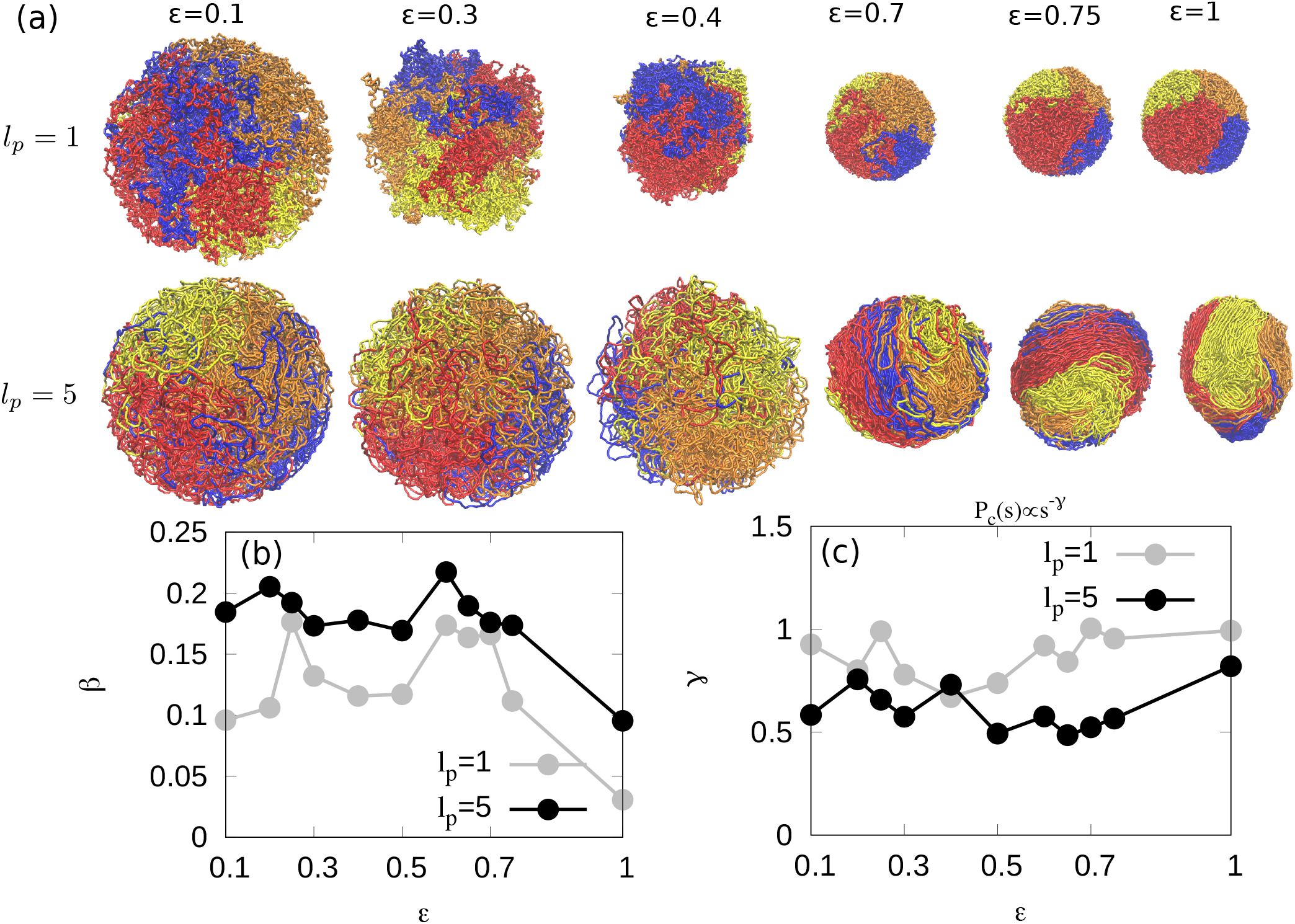
Mixing of four chains in small confinement volume (for a relatively large volume fraction of chains, *ϕ* = 0.1) varying the bead-bead interactions, *ϵ* : (a) Simulation snapshots of the mixing of four chains as a function of the interaction strength, *ϵ*, for persistence lengths *l*_*p*_ = 1 bead and *l*_*p*_ = 5 beads. (b) Scaling exponent *β* of the time dependence of the the chromosome mixing index. (c) Scaling exponent *γ* of the contact probability as a function of *ϵ* for *l*_*p*_ = 1 (gray color) and *l*_*p*_ = 5 (black color).

**FIG. A.6:**
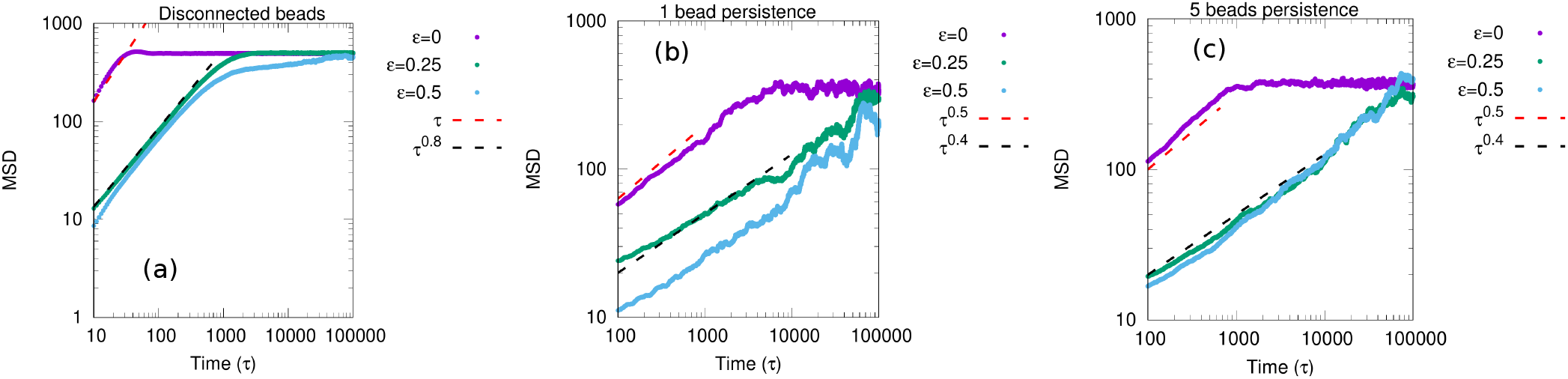
Mean-square displacement (MSD) of a bead (averaged over all the beads of the chain) confined chromosome chains: MSD are calculated from simulations for phantom chains (*ϵ* = 0), repulsive chains (*ϵ* = 0.25), and attractive chains (*ϵ* = 0.5) for a confinement volume equivalent to a bead volume fraction of *ϕ* = 0.4. Within this confinement, the MSD of the bead increases with time, and when its value reaches the square of the confinement radius, the MSD saturates at a constant value. (a) When beads are disconnected, for *ϵ* = 0, MSD increases linearly with time (MSD ~ *τ*) and for *ϵ* = 0.25 and 0.5, MSD ~ *τ* ^0.8^. These results demonstrate that for *ϵ* = 0, beads behave as independent diffusive particles, and sub-diffusion occurs when we introduce interaction (repulsion or attraction) between them. MSD of chain with a persistence length of (b) 1 bead and (c) 5 beads, for different interaction strength, *ϵ* are shown. For *ϵ* = 0, MSD ~ *τ* ^0.5^ is the result expected from Rouse chains [12]. For repulsive (*ϵ* = 0.25) and attractive (*ϵ* = 0.5) interactions, MSD ~ *τ* ^0.4^ [52].

**FIG. A.7:**
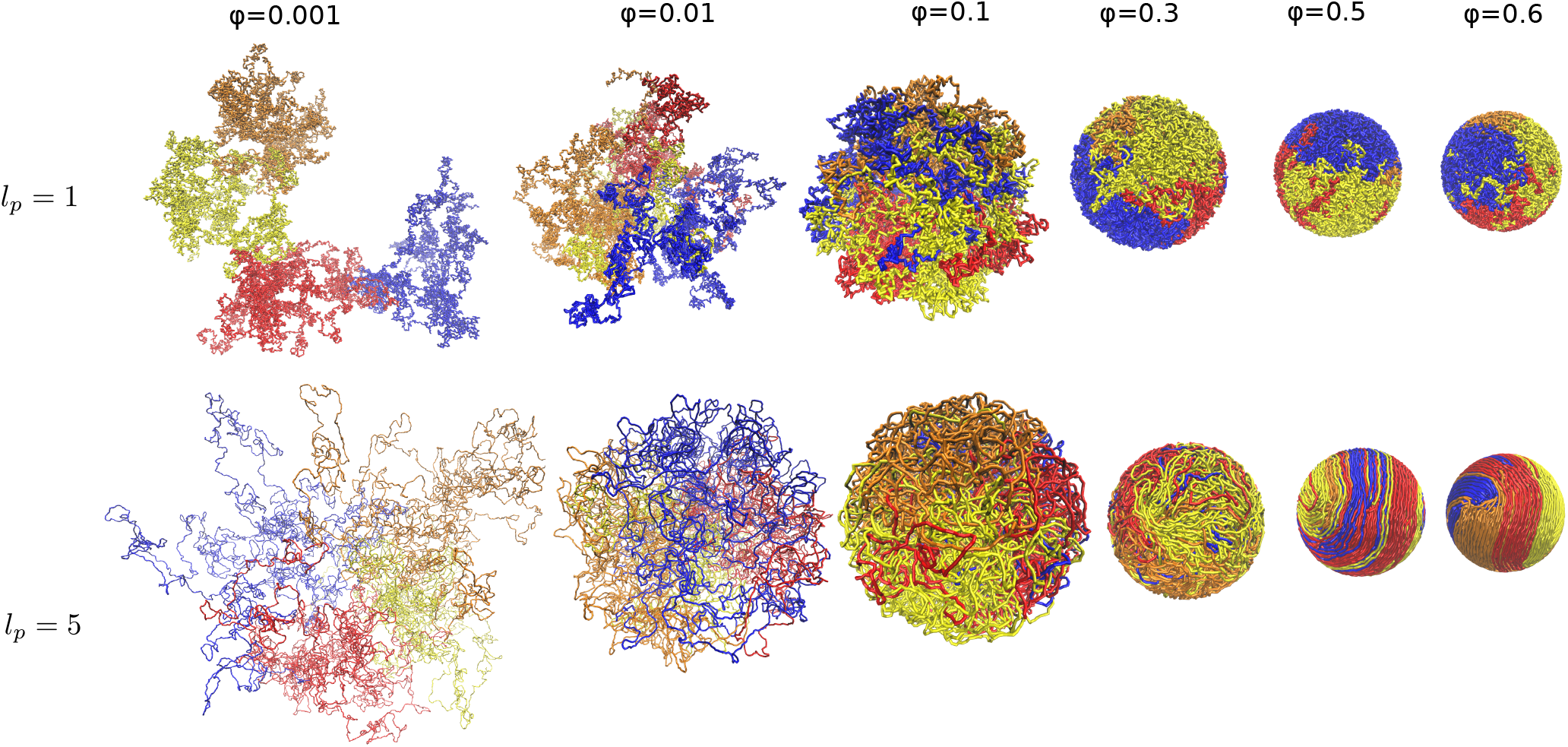
Mixing of four chains with weak attraction strength *ϵ* = 0.25 while varying the confinement volume (equivalent to fixing the volume fraction of chains *ϕ*): Simulation snapshots of the mixing of four chains as a function of volume fraction, *ϕ*, for persistence lengths *l*_*p*_ = 1 bead and *l*_*p*_ = 5 beads. Note that some snapshots were zoomed out because they were too large and would take up too much space if they were shown at their actual size.

